# Genome assembly of the C_4_ grass *Themeda triandra* confirms karyotype orthology with sorghum and reveals genetic variation across diverse environments

**DOI:** 10.64898/2026.03.18.712786

**Authors:** Jakob B. Butler, Jazmine L. Humphreys, Theodore Allnutt, Vinod K. Jacob, Lily Chen, Alejandro Correa-Lozano, Javier López-Jurado, Eloise Foo, Ian J. Wright, Steven M. Smith, Brian J. Atwell

## Abstract

*Themeda triandra* (tribe Andropogoneae) is a pan-continental grass occurring throughout Australia. Recognising its remarkable adaptive capacity, we assembled the *de novo* nuclear genome sequence of a diploid accession from eastern Australia, finding all ten chromosomes of the 755 Mb assembly highly syntenic with those of *Sorghum bicolor*. Genotyping and cytometry of accessions from across the climatic range of Australia show geographically structured lineages, with polyploids dominating in arid regions, apart from a notable diploid lineage from the arid north-west of Australia. A detailed comparison of this arid-zone lineage with two diploids from temperate south-eastern Australia through whole-genome resequencing revealed extensive putative copy number variation and polymorphisms, with changes in orthologs of known flowering regulation genes in *Sorghum* consistent with the environmental origin of *Themeda* accessions. Exploration of the extensive adaptation and resultant genetic diversity in *T. triandra* is expected to benefit grassland management programs and enable introgression of novel genes into Andropogoneae crops for climate resilience.

**SIGNIFICANCE STATEMENT:** *Themeda triandra* spans cool-temperate to tropical climates across three continents without speciating, making it a natural model for climate adaptation in C_4_ grasses. Its first *de novo* chromosome-scale genome reveals conserved karyotype orthology with sorghum, and population genomics identifies geographically structured lineages and candidate adaptive loci, providing a foundation for linking natural variation in a keystone grassland species to climate resilience in C_4_ crops.

## INTRODUCTION

The tribe Andropogoneae is a pan-tropical complex of 90 genera and 1,200 species of C_4_ grasses, many of which are key species in savanna ecosystems [1]. It also includes major crops such as maize, sorghum and sugarcane. Within the subtribe Anthistiriinae, *Themeda triandra* exemplifies the remarkable geographic breadth of the tribe, occurring naturally across south and south-east Asia, the Middle East, eastern and southern Africa, and Australasia [2–4]. Within Australasia, *T. triandra* (formerly *T. australis*) occurs naturally across every climate zone, extending from the northern tropics to the southern cool-temperate and maritime regions to latitudes of more than 40°S [5, 6].

*Themeda triandra* has a generally perennial habit in the field, with vegetative growth in spring culminating in sexual and aposporous flowering during warmer seasons when soil water is available [7]. However, populations in different climate zones exhibit extensive phenotypic variation; for instance, accessions collected from New Guinea to Tasmania have markedly different photoperiod responses [7, 8]. Even within accessions from SE Australia, morphological traits such as tiller number and flowering time vary markedly [6]. Variation in awn length occurs according to climate regime, with cool southern Australian accessions differing from those inland, which are in turn distinct from accessions from the arid north-west [9].

Common garden experiments have shown that certain characters are determined by the site-of-origin of different accessions, indicating that different populations are genetically distinct. For example, the timing of reproductive development among accessions, as estimated by the days from germination to emergence of inflorescences, was found to be related to the climate-of-origin [7]. More recently we have found that in two temperature-controlled treatments, morphological and physiological traits during vegetative development correlated weakly with the temperature and precipitation regimes at the site-of-origin of accessions [8]. When maxima were constrained to 30 °C, accessions originating in the wet sub-tropics took 1 - 2 weeks longer to flower under long days, in comparison to temperate accessions. When grown at sub-optimal maxima (20 °C), this difference was exacerbated, with the sub-tropical accessions delayed by an average of five weeks [8]. Thus, cool growing conditions unmasked a ‘time to flowering’ phenotype among different accessions and hence, a strong genotype × environment interaction.

The evolutionary origins of *T. triandra* and specifically, its spread to Australia in the past 1 - 2 million years [3], are critical considerations for understanding this divergence within the species. Phylogenetic analysis based on plastid and mitochondrial DNA sequencing of Australian and global accessions has suggested at least two independent colonisation events [3, 10]. Subsequent genetic fragmentation has been documented across SE Australia, where climate regimes and geographic separation have driven diversity in the ploidy and genotypes of 52 accessions [11]. Diversity between populations of *T*. *triandra* across SE Australia has been shaped by the strong selective forces of temperature and available water, together accounting for a quarter of genetic divergence in these populations [11].

Polyploidy is common in wild plant populations, often thought to influence rates of speciation and adaptation to stress [12]. Polyploids ranging up to octoploids in local populations of *T. triandra* throughout Australia have been reported [5, 11, 13]. This widespread polyploidy is consistent with the species’ ecological success across a vast range of climates and soil types [14]. Polyploid *T. triandra* populations are generally better adapted to stressful environments, exhibiting faster vegetative growth rates [15]. Furthermore, tetraploid populations from coastal New South Wales have reproductive advantages compared to diploids [14]. The co-existence of *T. triandra* populations that have admixtures with different ploidy levels raises more nuanced questions of adaptation [10].

Other mechanisms through which *T. triandra* may have adapted to such diverse environments include genome organisation, gene copy number and mutations affecting gene expression and function. Such genome-level phenomena are key targets for the transfer of traits from wild crop relatives to molecular breeding of superior crops [16] but first require extensive genetic characterisation. Genome assemblies have recently been reported for *T. triandra* accessions from The Philippines and Australia [10, 17], although only to the contig level *de novo*. Access to a complete *de novo* reference genome sequence is now essential for deeper investigation of the evolution of *T. triandra* and its application to cultivated species.

The aims of this study were to generate a chromosome-scale reference genome for a diploid *T. triandra* accession from eastern Australia, and to use this resource alongside whole-genome resequencing of accessions from contrasting climates to characterise copy number variation and sequence polymorphisms that distinguish divergent lineages. We further aimed to combine SNP genotyping and ploidy assessment across a broader sample set to reconstruct phylogenetic relationships among disparate Australian populations and examine patterns of radiation and genomic differentiation associated with climate.

## MATERIALS AND METHODS

### Themeda accession sampling and cultivation

Samples of *T. triandra* were selected from 20 different localities representing diverse climatic origins across the Australian continent, spanning mean annual temps from 10.9 - 26.9 °C and annual precipitation from 281 - 1198 mm (Supplementary Table 1, Figure 1). Accessions were sourced from suppliers including the Australian Pastures Genebank, Native Seeds Pty Ltd and Nindethana Seed Service Pty Ltd as well as personal collections by BJA and ACL. As personal collections were opportunistic, some populations were represented by multiple seeds from a single plant (Supplementary Table 1).

**Fig. 1.**
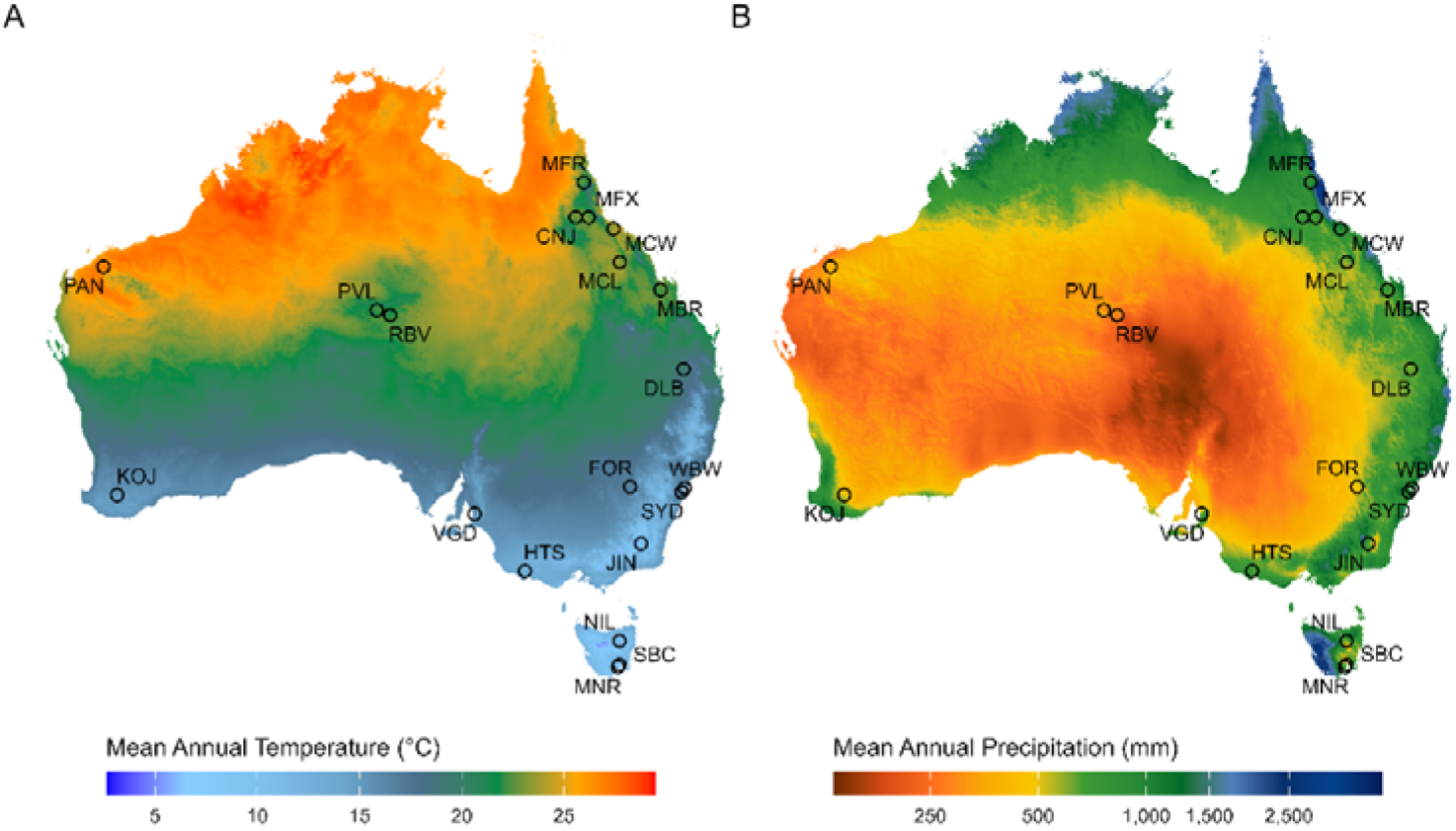
Geographical locations of *Themeda triandra* accessions used in this study. The maps of Australia are colour-coded according to A) mean annual temperature and B) mean annual precipitation, with variables obtained from gridded datasets from CHELSA [18] at a resolution of ∼1 km at the equator and representing the long-term interpolated average of locations between 1981 – 2010. The three letter notations are expanded upon in full in Supplementary Table 1.

Seeds were germinated in a controlled growth chamber (at 35/20 °C in a 12-h day/night regime, with a peak irradiance of 1000 μmol photons m^−2^ s^−1^ midway through the photoperiod) and transplanted to 4.45-L pots containing a 9:1 mixture of commercial potting mix (<30% fine sand, >70% pine bark; Australian Growing Solutions) and field soil (40:30:30 sandy clay loam). Plants were maintained in climate-controlled glasshouses under a 30/26 °C day/night temperature regime with a photoperiod of 14 h (natural light was supplemented when falling below 700 μmol photons m^−2^ s^−1^ with blue/red LED lighting, adding 600 μmol photons m^−2^ s^−1^).

For genomic analyses, leaf tissue was sampled from 3-month-old plants and immediately frozen in liquid nitrogen and stored at −80 °C until DNA/RNA extraction.

### Library construction and sequencing of reference individual

Leaf tissue samples of the “WBW” accession were sent to Biomolecular Resource Facility (Australian National University) for high molecular weight genomic DNA extraction and long-read library construction using a SMRTbell® prep kit 3.0, with BluePippin size selection used during library preparation. PacBIO-HiFi sequencing was undertaken using the Revio SMRT Cells platform, generating 96.7 Gbp of HiFi long reads with an expected coverage of the *Themeda triandra* genome of 114X, based on the 1C genome size of 0.84 Gbp for *T. triandra* estimated previously [10]. Further leaf tissue samples were also used to prepare a Hi-C library from cross-linked chromatin using an Arima Hi-C v2.0 protocol. This library was sequenced using Illumina NovaSeq 6000, producing 154.1 Gbp of Hi-C reads. RNA was extracted from five different tissues (leaf meristem, mature lamina, root tips, mature root and embryo) of the “WBW” accession using Kinnex full-length RNA kits, preserved in liquid nitrogen and sent to AGRF Australia (Melbourne) for long-read sequencing using PacBio ISOseqX, with 94.0 Gbp of transcript sequence generated.

### Genome assembly and annotation

PacBio HiFi reads were assembled using hifiasm v0.19.18 [19]. Contigs belonging to mitochondria and chloroplasts were identified by BLAST search of known sequences, and those contigs were extracted. After haplotigs were removed using purge_haplotigs v1.1.2 [20], genome heterozygosity was estimated from HiFi reads by counting canonical k-mers (k = 21) with Jellyfish v2.3 [21] and modelling the resulting frequency histogram with GenomeScope 2.0 [22]. The remaining contigs were then scaffolded using deduplicated Hi-C reads and YaHS v1.2a.1 [23]. The Hi-C contact map was visualized using Juicer v1.6 (https://github.com/aidenlab/juicer). Repetitive elements were discovered using the ENSEMBL [24] repeat annotator and soft-masked in the final assembly, while 74 scaffolds containing >95% repetitive DNA were discarded entirely, along with 4 scaffolds which contained contaminants as determined by NCBI Foreign Contamination Screen [25]. Genome (and annotation, see below) completeness was assessed using BUSCO (poales_odb10 library) [26] *via* compleasm v0.2.5 [27]. The occurrence of telomeric repeats were detected and annotated using tidk v0.2.41 [28], while genome-wide repeat content, including LTR retrotransposon families, was characterised in full using EDTA v2.2.2 [29].

Structural gene annotation was performed on the ENSEMBL soft-masked sequence using BRAKER3 v3.0.8 [30], using protein sequences from *Sorghum bicolor* obtained from NCBI (Taxonomy ID 4558) as external hints to guide gene model prediction and the aligned ISOseq transcript data for evidence of model support. Gene models with less than four residues in the translated product were discarded. Predicted protein-coding genes were functionally annotated using InterProScan v5.66-98.0 [31] and eggNOG-mapper v2.1.12 [32], with KEGG orthology determined using BlastKOALA v106.0 [33]. Annotation of rRNAs was performed using barrnap v0.9 (https://github.com/tseemann/barrnap).

To compare with published *T. triandra* assemblies (GenBank GCA_018135685.1, GCA_025603265.1), contiguity statistics were recalculated for all three assemblies from the GenBank sequences on a common basis (nuclear sequence only; contigs defined by splitting scaffolds at runs of ambiguous bases). Each assembly was then aligned to the WBW reference with minimap2 v2.1 [34], retaining primary alignments with MAPQ ≥5 and block length ≥1 kb, with genome coverage and gene recovery computed against the WBW annotation.

### Genome synteny with other Andropogoneae

Synteny with the *Sorghum bicolor* reference genome (BTx623 v3.0, [35]) was assessed using *nucmer* v3.1 [36], anchoring on maximal exact matches unique in *Sorghum*, with a minimum exact match length of 40 bp, a minimum cluster length of 200 bp, a maximum gap of 1,000 bp between adjacent matches within a cluster, and an alignment extension break length of 500 bp. Detected structural variation was quantified using *syri* v1.7.0 [37]. Given the high level of colinearity detected between these genomes, *T. triandra* chromosomes were named and oriented to maximise synteny with the *S. bicolor* genome.

To characterise patterns of gene family evolution, synteny, and sequence divergence between the major Andropogoneae crops and *Themeda*, multiple comparative analyses were performed using algorithms implemented in OrthoVenn3 v1.1.1 [38], comparing the gene content of *T. triandra* to that reported in the reference genomes of *S. bicolor* (BTx623 v3.0) and *Zea mays* (B73 v5.0, [39]). Orthologous gene clusters were identified using OrthoMCL [40], with an e-value threshold of 1 × 10^⁻²^ applied to define cluster membership. Gene family expansion and contraction along each lineage since divergence were inferred using CAFE5 [41]. Pairwise collinearity of genomic regions containing at least five orthologous genes was assessed among the three species using MCScanX [42]. In addition, protein sequences of the single-copy orthologous genes shared among species identified using OrthoMCL were aligned using MAFFT v7 [43], converted into codon alignments using PAL2NAL v14 [44], and pairwise rates of synonymous substitution (Ks) were calculated for each gene using the PAML v4.10.9 program codeml [45] to assess relative rates of sequence mutation and divergence.

### Sampling and genotyping using ddRAD sequencing

DNA was extracted from 103 accessions (with 40 samples duplicated) from 18 populations (Supplementary Table 1) using the CTAB method [46] with additional RNAse treatment. For genotyping *via* ddRAD sequencing, DNA was shipped to AGRF Australia. The DNA underwent PstI/HpyCH4IV restriction digest, insert size selection (280 - 375 bp) using BluePippin, library preparation and sequencing using NovaSeq X Plus (Illumina Inc., San Diego, CA, USA), with 633.5 Gbp of paired-end sequence generated and an average of 4.4 Gbp per sample. Using fastp v0.24.0 [47], reads were trimmed of poly-G artefacts, adapters were automatically detected and removed, a sliding-window quality trim (4 bp window, mean quality threshold Q20) was applied, and reads shorter than 75 bp after trimming were discarded. Reads were then demultiplexed using stacks v2.65 [48], specifying the options for cutsite and barcode rescue, removing reads with missing data and low quality, and ignoring the absence of the cut site on reverse reads due to a diversity issue on reverse reads based on the specific library construction [49]. Given this issue the first three bases of the reverse read were also trimmed using seqtk v1.3 (https://github.com/lh3/seqtk). Adapters were then removed using trimmomatic v0.39 [50], and all reads (both paired and orphaned) were processed and aligned to the *T. triandra* genome using bwa mem v0.7.17 [51] and samtools v1.16.1 [52]. As some accessions were duplicated, the samples with greater amounts of sequencing were retained. SNP calling was then performed using the mpileup and call tools from bcftools v1.15.1 [52], with a maximum per-sample depth of 1000x and a minimum mapping quality of 20. The resulting SNPs were then filtered iteratively; individuals with over 90% missing genotypes were removed, followed by removing SNPs that did not meet the following criteria: maximum missing proportion of 20%, a minimum depth of 5, and a minimum minor allele frequency of 0.05. Subsequently, a SNP pruning step was undertaken to randomly remove a SNP from pairs of SNPs in strong linkage disequilibrium with each other (r^2^ > 0.6) using bcftools prune.

### Whole genome resequencing of six accessions

Full genome resequencing was performed on six selected accessions (Supplementary Table 1). DNA was extracted from leaf samples of these accessions using the CTAB method [46] with RNase treatment and delivered to AGRF Australia for library preparation with Illumina DNA PCR-Free kits and sequencing, with 176.6 Gbp of paired end sequence generated using NovaSeq X Plus (Illumina Inc., San Diego, CA, USA), equating to approximately 39X coverage per sample. Reads were quality controlled using fastp [47] and aligned to the *T. triandra* reference genome using bwa mem. Automatic detection of potential DNA structural variation in these samples *via* disjunct alignment of reads was performed using manta v1.6.0 [53]. SNP calling was then performed against the *T. triandra* reference genome as with the ddRAD data. For subsequent analysis, SNPs called from the resequencing of one accession (KOJ-a) and two outgroup accessions from the Philippines (BioSample SAMN30450876) and China (BioSample SAMN08770639) were intersected with the filtered ddRAD SNP set, with a subsequent missing data filtering of 90% to reduce artefacts arising from combining sequencing technologies [54].

### Determining accession ploidy

Ploidy measurements were initially based on flow cytometry on select samples from across the range to validate results of the ddRAD data analysis (below). Approximately 0.5 cm² (or a ∼2 cm segment) of young leaf tissue was rapidly chopped for 60 s to fineness with a razor blade (replaced after 10 samples) in a petri dish containing 250 µl of Nuclei Extraction buffer supplied in the CyStain UV Precise P kit (Sysmex Partec; Order No. 05-5002). A further 250 µl of extraction buffer was added to yield ∼500 µl of suspension (plus debris). The suspension was pipetted through a 30 µm CellTrics® filter (Order No. 04-0042-2317) into a transparent polypropylene 3.5-ml Sysmex sample tube (Order No. 04-2000). Then, 2 ml of CyStain UV Precise P Staining buffer was added and samples were incubated at room temperature for 60 s. Flow measurements were performed by inserting the 3.5-ml tube into a CyFlow® Ploidy Analyser (Sysmex Partec) equipped with a UV LED suitable for DAPI excitation. The instrument was operated with 1× PBS as sheath fluid and data were acquired using CyView v2.4 software. A sheath fluid flush was performed between each sample.

A DNA control standard supplied by the manufacturer (Sysmex) was run at the beginning of each session as a quality control check. Reference *T. triandra* accessions used to assign ploidy were: 1) a diploid accession from Falmouth, Tasmania (APG 85441) and a 2) a tetraploid accession from Winderlup, near Mylor, South Australia (35.038 S 138.772 E) provided by Marne Dunin (pers. comm.). Reference extracts were used to align G /G peaks. Instrument settings, including gain and flow rate, were optimised using the diploid standard to position its G /G peak within the linear detection range; these settings were then maintained for all subsequent samples within that run, allowing ploidy levels (diploid, tetraploid or hexaploid) to be assigned based on the relative position of G /G peaks compared to the diploid standard. Histograms with a coefficient of variation (CV) of the primary G /G peak greater than 6% were excluded, and the corresponding samples were re-analysed.

Ploidy of the samples in the ddRAD data were also assessed by the proportion of heterozygous reads, following the methods of Ahrens, James [11], with samples shared between cytometry and genotyping methods used for validation. The read depth of each allele was extracted for all SNPs within all samples from the variant call format (VCF) file, and the relative frequency of each allele was calculated. A histogram was made for each sample, showing allele frequencies of all non-homozygous SNPs, and sample ploidy was assigned by visual analysis of the distribution. Assuming consistent read coverage, each heterozygous locus in a diploid individual should have a read-depth ratio of 1:1 for the reference and alternate allele, resulting in a normal distribution of allele frequencies with a single peak at a reference allele proportion of 0.5. Higher ploidy would have different distributions of allele frequencies at heterozygous loci, resulting in one additional peak for each step increase in ploidy (e.g. heterozygous loci in triploid accessions would have expected reference allele read-depth proportions of 0.33 and 0.66, tetraploid accessions 0.25, 0.5 and 0.75, etc). After assessing SNPs of reads depths 5 - 400, only SNPs with a read depth > 200 were used, as this gave the greatest peak resolution in this data. Determining the ploidy of the resequenced accessions intersected with these data required reducing the depth threshold to 50 to achieve similar resolution.

### Phylogenetic analysis

To investigate genetic relationships and diversity among accessions, a maximum likelihood phylogeny was created using a workflow adapted from Severn-Ellis, Scheben [55]. Genotypes were re-called in a ploidy-aware manner using the allele-ratio assignments with FreeBayes v1.3.6 [56], retaining genotype qualities and up to four alleles per site. Calling at the appropriate ploidy avoids low-dosage alleles in polyploids being called as homozygous. The filtered VCF was converted to PHYLIP format using vcf2phylip v2.0 [55], which encodes heterozygous genotypes as IUPAC ambiguity codes, so allele dosage is not retained and the topology reflects allele sharing among accessions rather than ploidy. Maximum likelihood phylogenetic inference was performed using RAxML v8.2.12 [57] using the ASC_GTRCAT model and the Lewis ascertainment bias correction with no among-site rate heterogeneity, with 100 rapid bootstrap replicates to assess node support. The phylogeny was projected into geographic and climatic space using the R package phytools v2.5-2 [58].

Several samples were removed from analysis due to mislabelling of accessions identifiable from the genotyping data/phylogeny (Supplementary Table 1). In total, 14 samples were identified as mislabelled and removed due to unexpected population grouping within single-family clades. Specifically, in clades with very short branch lengths (indicating samples of seed from a single plant), mixed populations were identified as problematic, with samples removed based on geographic disagreement with the population in the nearest clade.

### Comparison of CNV and SNP impacts in resequenced accessions

To detect putative copy number variations (CNV) among contrasted diploid accessions, changes in resequencing depth were examined using CNVkit v0.9.11 [59]. Normalised read depth profiles for accessions from Tasmania (SBC), Sydney, (SYD) and Pannawonica (PAN) were generated relative to a flat reference, providing a common baseline for estimating copy number across the genome, although with the caveat that allelic divergence between accessions and the reference individual may reduce mapping in divergent regions. Overall heterozygosity of these accessions was assessed independent of the reference genome by counting canonical k-mers (k = 21) in the trimmed reads with Jellyfish and Genomescope2 to evaluate the potential extent of this divergence. Pairwise comparisons between accessions (PAN–SBC, PAN–SYD and SBC–SYD) were then performed to identify differences in gene-level sequencing depth. Each 5 kb segment of the chromosomes that intersected with a gene was collated (discarding those genes that spanned two segments with opposite signs in fold change). Log2 fold changes in depth were then calculated for each gene, with values > 1 conservatively classified as copy number gains, values < −1 classified as copy number losses, and genes with mean read depth ≤ 5 in one accession and > 5 in the other interpreted as putative deletions. As heterozygous copy number changes produce log2 depth ratios of approximately −1 (hemizygous loss) and +0.58 (hemizygous gain), these thresholds are conservative and preferentially recover homozygous or higher-order copy number differences. To assess for potential functional differences between these accessions, the impact of high-quality SNPs (depth > 5, quality score > 20) on annotated genes was determined using SnpEff v5.2 [60]. Genes containing SNPs with putative high impact on protein function (modification of start/stop codons, premature truncation and frameshifts) that differed between the three diploid contrasts were identified. All gene lists were tested for enrichment of GO terms using the R package topGO v2.64.0 [61] with Fishers exact test.

## RESULTS

### 1. Sequencing and assembly of a reference genome

We assembled a collection of *T. triandra* accessions from established seedbanks, herbarium collections and field sampling, spanning a wide range of Australian climatic regions (Supplementary Table 1, Figure 1). From this collection, a diploid accession from Brisbane Water National Park in Wondabyne (WBW) on the warm-temperate south-eastern coast was selected for reference genome assembly. We generated a total of 96.7 Gbp (114X) of PacBio HiFi reads and 154.1 Gbp of Hi-C sequencing for the assembly of the *T. triandra* WBW genome. The assembled genome was resolved into 10 chromosomes and 12 unassigned contigs with a total size of 755 Mb (Figure 2, Supplementary Table 2), closely matching the k-mer-based genome size estimation (733 Mb) indicating that allelic haplotigs were effectively removed to create a single haplotype. This assembly contains 75% of the total expected telomeres and a contig N50 of 61.7 Mb, indicating a highly contiguous assembly had been achieved, supported by a BUSCO completeness score of 99.74% (Supplementary Table 2). Analysis of k-mers indicated that the WBW reference individual was highly heterozygous (1.67%), and approximately 173 Mb of the genome was represented by k-mers occurring at >10,000X coverage, reflecting abundant high-copy satellite and retrotransposon sequence. Correspondingly, a total of 439 Mb of repetitive sequence motifs covering 58% of the genome were annotated, consisting overwhelmingly of LTR-retrotransposons (Supplementary Table 3). Genome annotation based on *S. bicolor* protein hints and 94 Gbp of Iso-seq data predicted 44,156 protein coding genes, along with 825 ribosomal RNA genes (5S = 163; 5.8S = 204; 18S = 222; 28S = 236) which were localised to two main clusters (chromosomes 3 and 9). Overall annotation completeness lagged the overall genome, with BUSCO completeness estimated at 74.6% (Supplementary Table 2). Chromosomes were numbered and oriented to match *Sorghum bicolor*, reflecting the strong chromosomal synteny between the two genomes described below. Compared with the two previously published *T. triandra* assemblies, both generated from single long-read datasets, this assembly is one to four orders of magnitude more contiguous (contig N50 61.7 Mb, against 601 kb and 13.4 kb; Supplementary Table 4). Whole-genome alignment showed both to be broadly collinear with the present assembly but to recover only 64.0% and 40.8% of it (71.3% combined), leaving 17.4% of annotated genes and 62.9 Mb in regions ≥50 kb unrepresented in either previous genome (Supplementary Figure 1).

**Fig. 2.**
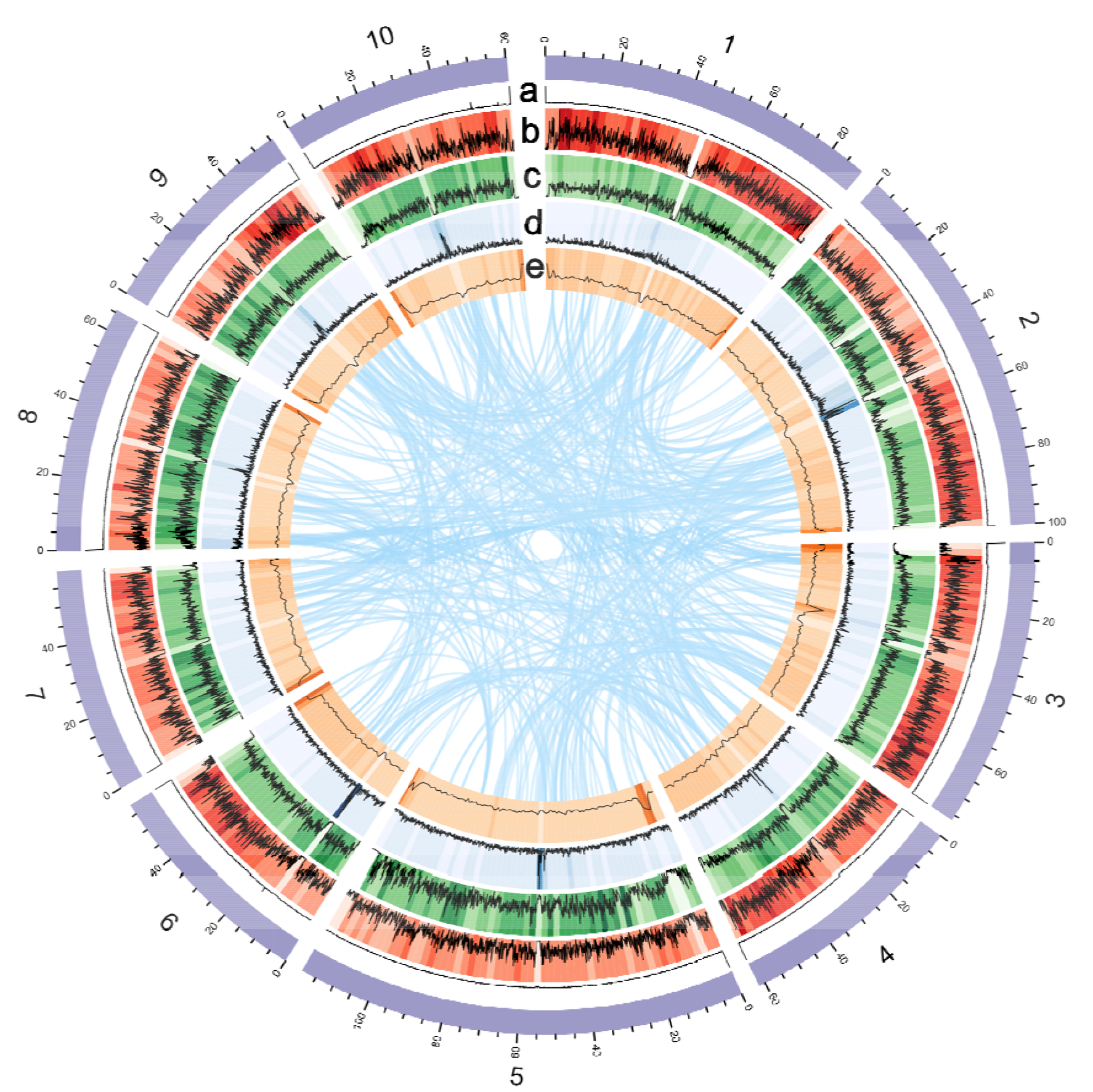
The genome of *Themeda triandra*. Outer ring of Circos plot shows the length of the 10 pseudomolecules (Mb). Inner data tracks are quantified across 100kb (line) and 2Mb (heatmap) windows: a) Number of telomeric repeats (0 – 500+); b) Gene density (0 - 19 per 100kb, 1 - 207 per 2Mb); c) SNP density (0 – 12,309 per 100kb, 239 – 107,360 per 2Mb); d) Repeat element density (0 - 187 per 100kb, 17 – 1,437 per 2Mb); e) %GC content (42 – 54% per 100kb, 42 – 53% per 2Mb). Link tracks indicate regions of strong gene synteny between chromosomes (that span ≥ 15 genes and ≥ 10 gene/Mb).

To complement the reference genome, whole-genome resequencing was undertaken for six accessions from different climatic regions around Australia. These included the hot arid north-west (PAN), the Mediterranean south-west (KOJ), the cool-temperate south-east (SBC and NIL), the warm-temperate south-east (SYD) and an alpine accession (JIN) (Figure 1). Resequencing enabled high-resolution SNP discovery and the identification of genomic regions with notable patterns of variation (Figure 2). Sliding-window analyses revealed multiple intervals exhibiting elevated or reduced SNP density, consistent with regions of suppressed recombination or localised haplotype divergence among lineages. When combined with both %GC content and gene density, other notable patterns were also distinguishable. The most obvious was the position of centromeric repeat regions, characterized by increased repetitive content and a reduction in %GC, gene density and SNP density. Likewise, the major cluster of rRNA genes on chromosome 3 was large enough to have a visible effect on genome composition (low gene, SNP and repeat density along with increased %GC, Figure 2). Detection of putative structural variants (SVs) comprising insertions, deletions, and rearrangements further supported this heterogeneity, with clusters of putative SVs concentrated in specific genomic regions (Supplementary Figure 2).

### 2. *Comparisons of* T. triandra, Sorghum bicolor *and* Zea mays

To provide a framework for transferring functional gene annotations from well-characterised crops to *T. triandra*, we compared its genome to those of *Sorghum bicolor* and *Zea mays*, the most extensively annotated and functionally studied Andropogoneae genomes. *Themeda* diverged from *Sorghum* ∼12 Mya, and this lineage split from *Zea* ∼15 Mya [3]. This makes *Sorghum* the closest well-resourced reference for inferring gene orthology, with maize serving as an outgroup. Whole-genome alignment revealed strong chromosomal synteny between *T. triandra* and *S. bicolor*, with each of the ten *T. triandra* chromosomes aligned predominantly to a corresponding *S. bicolor* chromosome, confirming a largely conserved macrostructure and implying orthologous karyotypes (Supplementary Figure 3). Across the genome, 6,725 high-confidence alignment blocks (>1 kb, ≥90% identity) were detected, with most alignments showing unique 1:1 correspondence, and a minority aligning to multiple locations consistent with segmental duplications. Despite this extensive synteny, many small intrachromosomal inversions were detectable, constituting approximately 94 Mb (12%) of the genome. These inversions were evident in alignment dotplots as clear diagonal breaks and reversals in local synteny (Supplementary Figure 3). The lack of inter-chromosomal translocations was notable, suggesting that genome evolution has proceeded *via* local structural changes rather than substantial chromosome rearrangement. Orthologous gene synteny also showed strong collinearity between *T. triandra* and *S. bicolor*, although approximately one-third of all syntenic gene blocks were inverted, as would be expected based on the established genome synteny (Figure 3a). Several *T. triandra* regions also show homology to more than one chromosome, particularly in *S. bicolor*, suggesting historical inter-chromosomal duplication of genic regions that have been maintained in both genera (Figure 3a). Some duplicated blocks were also detectable between *T. triandra* and *Z. mays,* although overall synteny was lower, as expected given its more ancient divergence and the maize-lineage specific whole genome duplication [62]. These patterns of synteny reveal substantial genome reorganisation after the divergence of *Zea* from the *Sorghum*/*Themeda* precursor, and that the broad reorganized state was subsequently maintained between these two genera.

**Fig. 3.**
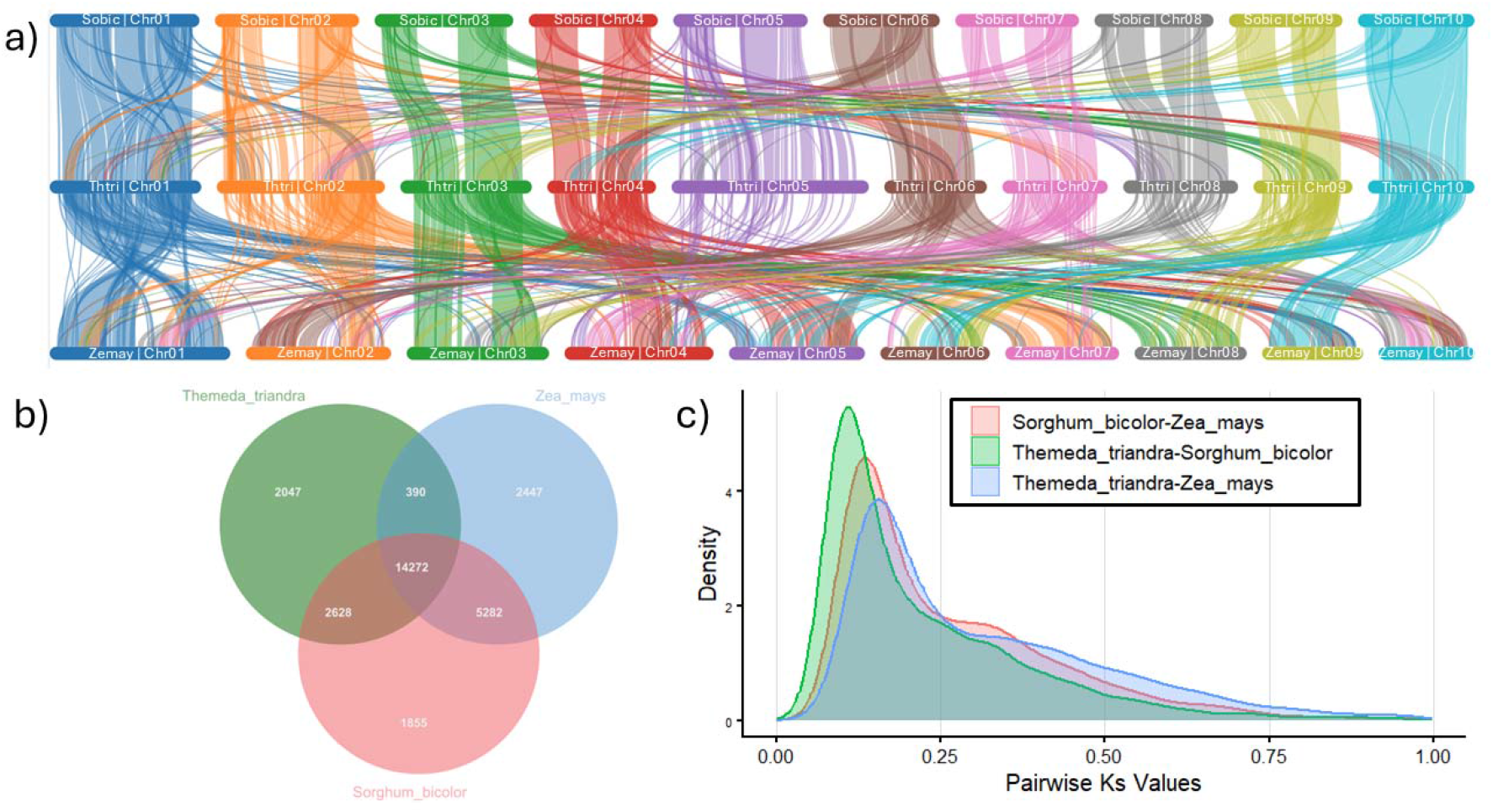
Comparison of orthologous genes between *T. triandra*, *S. bicolor* and *Z. mays.* A) Synteny of orthologous genes from *T. triandra* (centre) to *S. bicolor* (above) and *Z. mays* (b*e*low). Each link represents at least five genes in a collinear arrangement between the genomes compared. The central parts of some *S. bicolor* chromosomes have very few annotated genes, resulting in no visible links. B) Number of unique and orthologous gene families in *T. triandra*, *S. bicolor* and *Z*. *mays.* C) Distribution of synonymous mutation rate (Ks) values of 6,345 single-copy orthologues when compared pairwise between *T. triandra*, *S. bicolor* and *Z. mays*. A peak in Ks values indicates the most common number of neutral mutations found in these genes between the compared taxa, which can be linked to estimated divergence time and mutation rate.

Gene family analysis revealed 14,272 orthologous families (49%) conserved across all three lineages, with a further 2,047 families (7%) present in *T. triandra* but absent from *S. bicolor* and *Z. mays* (Figure 3b). The number of shared gene families varied between species pairs; while *T. triandra* shared 2,628 families exclusively with *S. bicolor*, it only shared 390 gene families exclusively with *Z. mays*, consistent with its more recent divergence time and greater synteny with the former. *Sorghum bicolor* and *Z. mays* nonetheless shared more gene families with each other (19,554) than either did with *T. triandra*, with 5,282 (18%) of those families absent from *T. triandra*. These families were not enriched for any particular function. The lower completeness of the new *T. triandra* annotation relative to the long-curated crop references (Supplementary Table 2) will inflate this figure (and the apparent contractions), but the 2,047 families recovered in *T. triandra* but not in *S. bicolor* or *Z. mays* were observed directly and cannot be an artefact of incomplete annotation. *Themeda g*ene families that were expanded, contracted or absent in *Sorghum/Zea* were each enriched for many of the same broad functional categories, including protein phosphorylation, ubiquitin-dependent protein catabolism, metal ion binding and response to oxidative stress (Supplementary Table 5a–c); 17 of the 39 GO terms enriched among expanded families were also enriched among contracted families. These overlapping enrichments are characteristic of large, dynamically evolving multigene families such as receptor-like kinases.

Pairwise synonymous substitution rates (Ks) calculated from 6,345 single-copy orthologues further supported these relationships. The Ks distributions showed peaks at 0.109 for *S. bicolor* – *T. triandra*, 0.134 for *S. bicolor* – *Z. mays* and 0.154 for *T. triandra* – *Z. mays* (Figure 3c). Assuming a *T. triandra – S. bicolor* divergence of ∼12 Mya and their split from *Zea* at ∼15 Mya [3] this corresponds to an estimated neutral mutation rate of 4.39 - 5.04 × 10⁻ substitutions per site per year. Notably, all three comparisons also showed a minor shared peak at Ks ≈ 0.33, reflecting a set of genes in the three species that have accumulated more neutral mutations than expected. Using the average mutation rate (4.66 × 10⁻□), this peak dates to ∼35.4 Mya. One explanation for this pattern could be ancient duplication followed by differential gene loss, consistent with the pattern of historic inter-chromosomal duplication. Significantly, only 2% of these genes (11 of 556) were located on different chromosomes in *T. triandra* and *S. bicolor*, instead suggesting that the 35.4 Mya peak reflects genes with deeper ancestral history rather than remnants of genome duplication.

### 3. *Biogeography, ploidy and phylogeny of Australian* T. triandra

Despite their genomic similarity, *S. bicolor* is largely adapted to semi-arid tropical and subtropical regions under irrigation, while *T. triandra* inhabits climates ranging from the hot, arid north-west of Australia to the cool-temperate southern regions. To understand the genetic contribution to this climatic versatility and the relationships between these distinct populations, we sampled *T. triandra* accessions selected from disparate sites throughout the species’ Australian range. Specifically, this panel of contrasting accessions originated from cool to hot regimes (mean annual temperatures 10.9 – 26.9 °C) and dry to wet regimes (annual precipitation 281 – 1198 mm) (Figure 1, Supplementary Table 1). Genotyping by ddRAD sequencing generated 50,395 filtered SNPs across the *T. triandra* genome.

Phylogenetic reconstruction showed that genetic relationships were broadly consistent with geographic distance between the 19 sampled populations (Figure 4a,b). Ploidy levels, determined by heterozygous allele ratios and validated with flow cytometry (Supplementary Figure 4; Supplementary Figure 5), ranged from diploid to hexaploid. While most inland arid environments were occupied by polyploids, diploid populations were not restricted to benign environments (SBC, NIL, MNR), also occurring in the hottest and most arid north-west (PAN) of the continent (Figure 4c). These diploid accessions were highly divergent from those sampled from the south-eastern region of the continent (Figure 4a), indicating substantial genetic differentiation that was independent of ploidy.

**Fig. 4.**
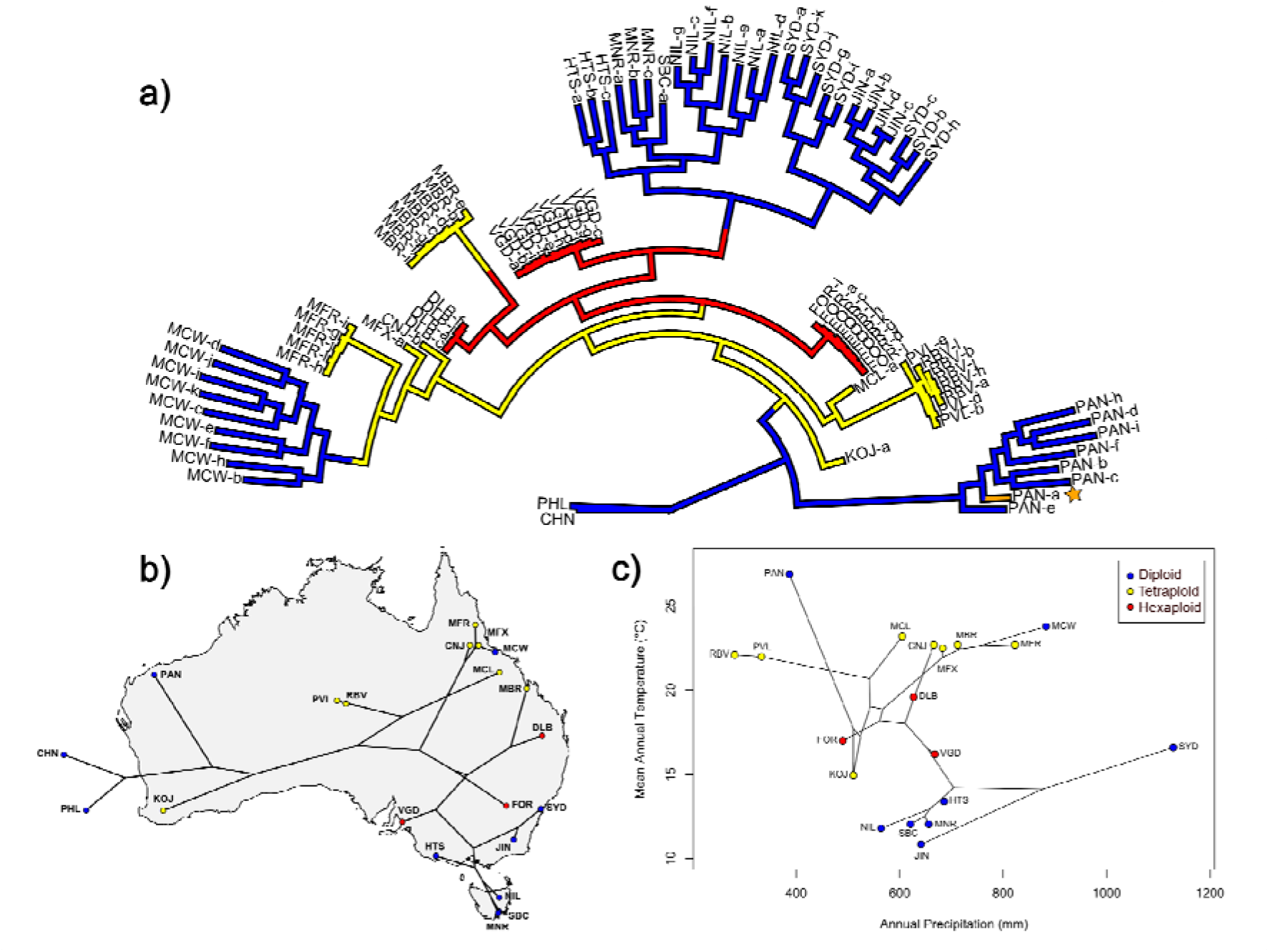
Nuclear phylogeny of *Themeda triandra* in Australia. A) Maximum likelihood radial phylogram of *T. triandra* populations across its Australian distribution. Ploidy of samples are indicated by tip colour, with 2n=blue, 3n=orange (highlighted with star icon), 4n=yellow, 6n=red. B) Maximum likelihood phylogram (with population clades collapsed) projected onto Australia based on coordinates of collection. Accessions PHI (Philippines) and CHN (China) as outgroups are placed outside of Australia. C) Phylomorphospace projection of phylogeny based on mean annual temperature and annual rainfall (mm).

Evidence of mixed-ploidy populations was observed at several sites: for example, genotyping revealed a triploid individual at Pannawonica (Figure 4a), likely arising from hybridisation between local diploids and neighbouring tetraploids (or meiosis error), and cytometric assessment at MFX indicated a mixture of diploid and tetraploid plants (Supplementary Figure 5). Ploidy was assigned independently of the phylogeny and mapped onto its tips (see Methods), so ploidy differences between neighbouring tips are not evidence of transitions along branches. Where a lower-ploidy accession sits among higher-ploidy relatives, the most likely explanations are that a single plant was sampled from a mixed-ploidy population or that independent colonisations of related lineages are being resolved together [3].

### 4. Genetic variation in diploids from climate extremes

To characterise the genetic divergence between diploid accessions from contrasting climates, SNPs from the resequencing of three diploid accessions (PAN, SYD, SBC) were annotated for their functional impact using SnpEff [60]. Across three pairwise comparisons, 11,159 unique genes carried high-impact variants predicted to strongly affect protein function, including modification of start/stop codons, premature truncation and frameshifts. 10,860 of these gene variants differed between either PAN-SYD or PAN-SBC, while only 5,675 differed between SYD-SBC, reflecting the higher divergence of the arid PAN lineage from the eastern diploids. Genes carrying these mutations were enriched for the GO term related to nucleic acid catalytic activity, but after correcting for multiple testing this enrichment was only significant in the SYD vs PAN comparison (Figure 5, Supplementary Table 6). Gene copy number variation between these accessions was also assessed using sequencing depth profiles with CNVkit [59], with 17,001 genes exhibiting putative CNV signals. 16,008 of these CNV signal differences were again present in the PAN-SYD or PAN-SBC comparisons compared to 6,550 in the SYD-SBC comparison. These figures may be inflated in part by PAN’s greater divergence from the reference reducing short-read mapping rather than reflecting true copy number loss, although the three accessions had comparable heterozygosity (1.58–1.90%, versus 1.67% for the reference individual), indicating similar within-genome diversity and hence a reference-divergence effect rather than a sample-specific one. Genes with putative copy number reduction in the PAN accession were significantly enriched for translation-related functions, including translational repression, regulation, initiation, and RNA binding activity, even after false discovery rate correction (Figure 5). Additional functional differences were identified across all three pairwise comparisons, with distinct sets of metabolic and transport-related functions enriched in each of the three accessions (Supplementary Figure 6, Supplementary Table 6).

**Fig. 5.**
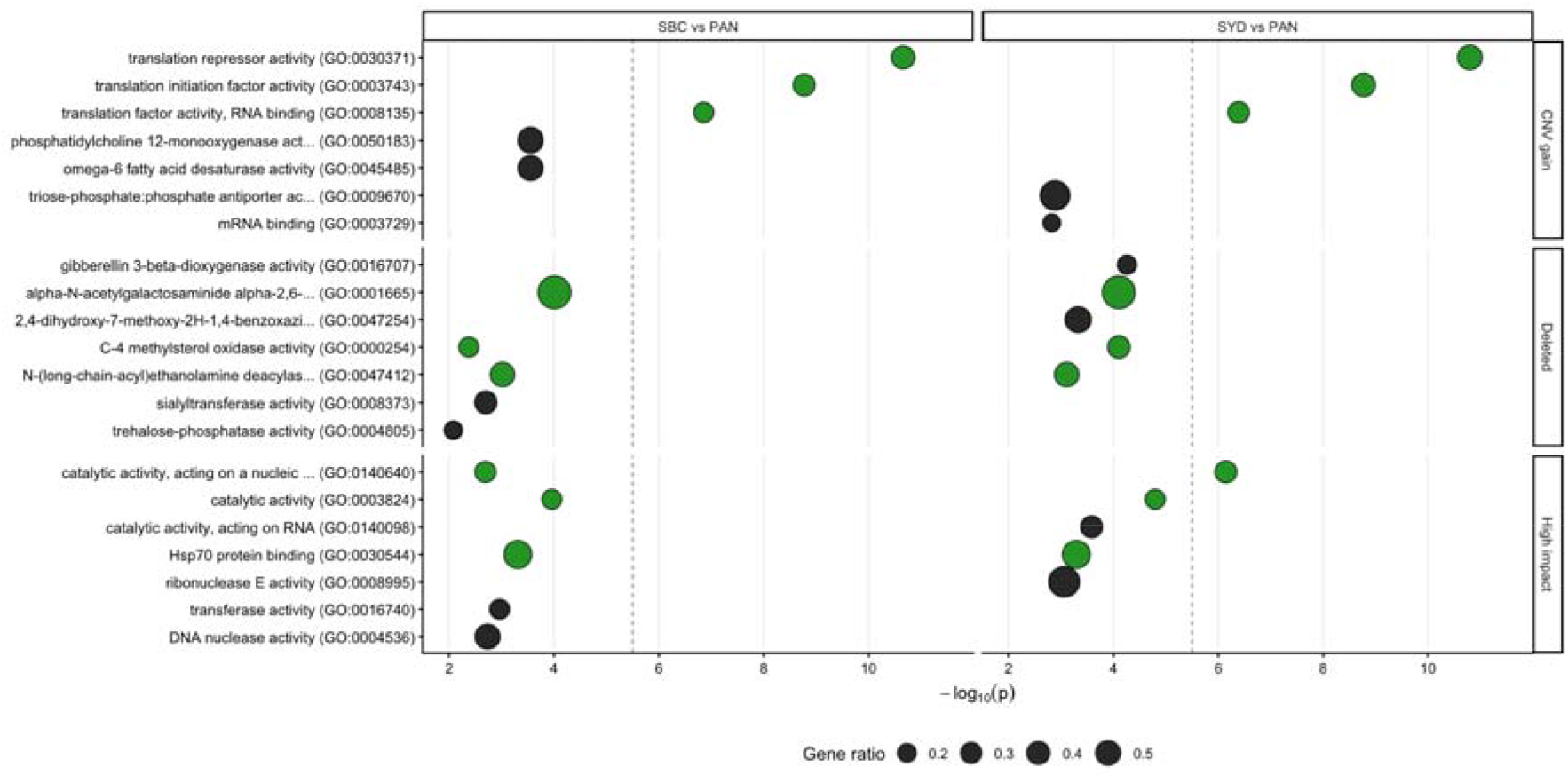
Top five enriched Gene Ontology (GO) terms of genes involved in putative copy number gain, deletions and housing high-impact mutations when comparing three diploid accessions of *T. triandra* from different environmental extremes. Accession names are described in Supplementary Table 1. The size of the bubble indicates what proportion of the entire set of genes annotated with that GO term are present in the set associated with putative copy number variation (CNV). Bubbles coloured green indicate GO terms significant in both comparisons. The vertical dotted line indicates the threshold after which P-values remain significant after correction for multiple testing using false discovery rate.

Previous work has shown that the *T. triandra* accessions in this sample originating from hot environments exhibit substantially delayed flowering at low growth temperatures compared with accessions from colder environments [8]. To investigate the genetic basis of this difference, variation in genes associated with flowering in these three diploid accessions representing environmental extremes was examined. Candidate genes were identified based on homology with known *S. bicolor* flowering genes [63]. Distinct patterns of functional and copy number variation in flowering-related genes were observed among all three accessions (Table 1). In the SBC accession from cool-temperate Tasmania, changes relative to the other accessions were concentrated around putative regulators of *FLOWERING LOCUS C* (*FLC*), a MADS-box transcription factor gene that acts as a potent repressor of flowering, specifically acting on *FLOWERING LOCUS T (FT)* floral promoter genes [64, 65]. These changes included copy number reduction in both a positive and negative regulator of *FLC*, as well as a high-impact mutation in the vernalisation-responsive gene *SbVRN1*, which also represses *FLC* activity. The SYD accession showed a partially overlapping but distinct set of changes relative to PAN, also centred on *FLC* regulation, with copy number increases in a positive regulator of *FLC* alongside the florigen *FT* gene and a negative regulator of *FLC*. In contrast, the PAN accession harboured high-impact mutations across a much broader set of flowering genes, spanning signalling, photoperiod response, and floral transition pathways, together with copy number reductions in several genes that promote flowering (Table 1). Assuming the function of these genes has been maintained since divergence from *Sorghum*, these patterns collectively suggest that populations from the hottest climate regime have undergone more extensive diversification of different genetic pathways interacting with flowering, whereas the colder eastern and south-eastern populations have evolved more targeted genetic changes focused on *FLC*-mediated repression of flowering. While background genome-wide differentiation between these accessions is high (see previous paragraph), the putative flowering genes were less often affected by CNV or high-impact variants compared to the genome-wide background (28/101 *vs* 23,691/44,156; Fisher’s exact OR 0.331, p = 1.90 × 10⁻), as expected for a conserved, functionally constrained gene set. Regardless, these candidate loci are consistent with, though not direct evidence of, climate-driven regulation of flowering-time.

**Table 1.**
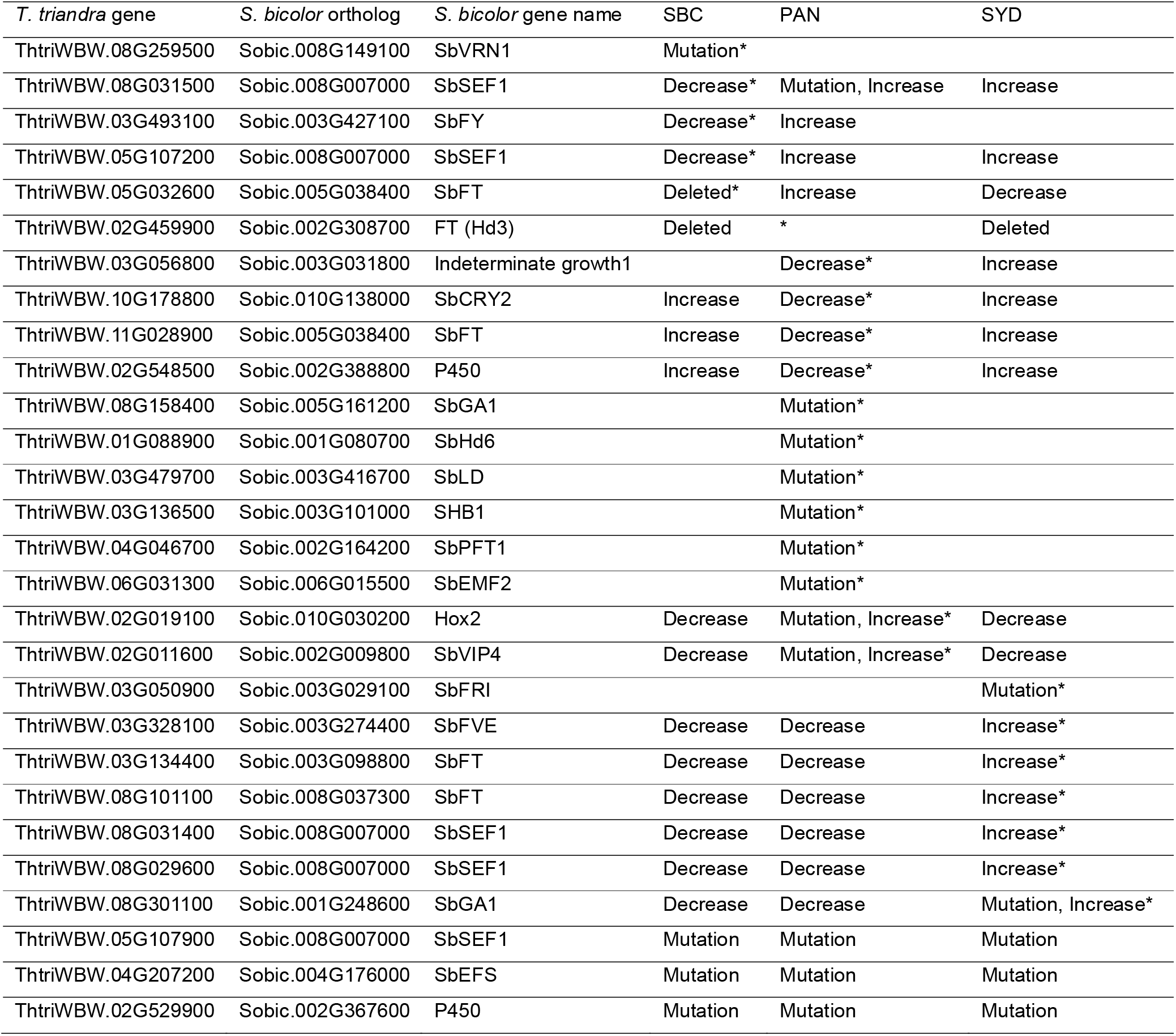
Copy number variation and high-impact mutations in *Sorghum* flowering orthologs across three diploid accessions of *T. triandra.* Increase: genes with putative copy number gain (log_2_fold change in sequencing depth > 1); Decrease: genes with putative copy number loss (log_2_ fold change in sequencing depth < −1); Deleted: genes with < 5 reads coverage; Mutation: genes with SNPs predicted to cause high impact changes to protein structure. Copy number variation is based on pairwise comparison between accessions. Blank cells indicate no difference meeting the above criteria between that accession and the others. Groups of classifications indicated by * highlight relevant accession patterns discussed in the main text. Accession names are detailed in Supplementary Table 1.

| *T. triandra* gene | *S. bicolor* ortholog | *S. bicolor* gene name | SBC | PAN | SYD |
| --- | --- | --- | --- | --- | --- |
| ThtriWBW.08G259500 | Sobic.008G149100 | SbVRN1 | Mutation* |  |  |
| ThtriWBW.08G031500 | Sobic.008G007000 | SbSEF1 | Decrease* | Mutation, Increase | Increase |
| ThtriWBW.03G493100 | Sobic.003G427100 | SbFY | Decrease* | Increase |  |
| ThtriWBW.05G107200 | Sobic.008G007000 | SbSEF1 | Decrease* | Increase | Increase |
| ThtriWBW.05G032600 | Sobic.005G038400 | SbFT | Deleted* | Increase | Decrease |
| ThtriWBW.02G459900 | Sobic.002G308700 | FT (Hd3) | Deleted | * | Deleted |
| ThtriWBW.03G056800 | Sobic.003G031800 | Indeterminate growth1 |  | Decrease* | Increase |
| ThtriWBW.10G178800 | Sobic.010G138000 | SbCRY2 | Increase | Decrease* | Increase |
| ThtriWBW.11G028900 | Sobic.005G038400 | SbFT | Increase | Decrease* | Increase |
| ThtriWBW.02G548500 | Sobic.002G388800 | P450 | Increase | Decrease* | Increase |
| ThtriWBW.08G158400 | Sobic.005G161200 | SbGA1 |  | Mutation* |  |
| ThtriWBW.01G088900 | Sobic.001G080700 | SbHd6 |  | Mutation* |  |
| ThtriWBW.03G479700 | Sobic.003G416700 | SbLD |  | Mutation* |  |
| ThtriWBW.03G136500 | Sobic.003G101000 | SHB1 |  | Mutation* |  |
| ThtriWBW.04G046700 | Sobic.002G164200 | SbPFT1 |  | Mutation* |  |
| ThtriWBW.06G031300 | Sobic.006G015500 | SbEMF2 |  | Mutation* |  |
| ThtriWBW.02G019100 | Sobic.010G030200 | Hox2 | Decrease | Mutation, Increase* | Decrease |
| ThtriWBW.02G011600 | Sobic.002G009800 | SbVIP4 | Decrease | Mutation, Increase* | Decrease |
| ThtriWBW.03G050900 | Sobic.003G029100 | SbFRI |  |  | Mutation* |
| ThtriWBW.03G328100 | Sobic.003G274400 | SbFVE | Decrease | Decrease | Increase* |
| ThtriWBW.03G134400 | Sobic.003G098800 | SbFT | Decrease | Decrease | Increase* |
| ThtriWBW.08G101100 | Sobic.008G037300 | SbFT | Decrease | Decrease | Increase* |
| ThtriWBW.08G031400 | Sobic.008G007000 | SbSEF1 | Decrease | Decrease | Increase* |
| ThtriWBW.08G029600 | Sobic.008G007000 | SbSEF1 | Decrease | Decrease | Increase* |
| ThtriWBW.08G301100 | Sobic.001G248600 | SbGA1 | Decrease | Decrease | Mutation, Increase* |
| ThtriWBW.05G107900 | Sobic.008G007000 | SbSEF1 | Mutation | Mutation | Mutation |
| ThtriWBW.04G207200 | Sobic.004G176000 | SbEFS | Mutation | Mutation | Mutation |
| ThtriWBW.02G529900 | Sobic.002G367600 | P450 | Mutation | Mutation | Mutation |

## DISCUSSION

The assembly of a chromosome-scale reference genome for *T. triandra,* combined with resequencing of accessions from contrasting environments, provides the comprehensive genomic framework required to investigate the success of this species across its remarkably diverse Australian range and beyond to Asia and Africa. The high degree of chromosomal synteny with *S*. *bicolor* opens the potential for translation of *T. triandra* genetics to improve major crop species. In addition, the genetic variation identified between accessions of *T. triandra* from contrasting biomes is a valuable resource for identifying genes and pathways required for tolerance to future climate regimes. Together, these make *T. triandra* a powerful resource for both understanding natural adaptation and informing crop resilience in the Andropogoneae.

The karyotype orthology between *T. triandra* and *S. bicolor,* with largely conserved gene order across all ten chromosomes, means that genes and regulatory regions in *T. triandra* can be mapped directly to their *Sorghum* orthologues. Despite approximately 12 million years of divergence, genome evolution between these lineages has proceeded primarily through intrachromosomal inversions rather than large-scale chromosome rearrangements, preserving this orthologous relationship. This finding is particularly relevant because cultivated sorghum has lost an estimated 14 - 35% of its wild progenitor’s genetic diversity through domestication [66–68], with modern breeding further narrowing the gene pool [69]. As a wild species spanning environments of unparalleled diversity, *T. triandra* retains adapted variation that has likely been lost in cultivated sorghum. While wild sorghum is clearly the most immediate source of new variation for sorghum breeding, the geographic and climatic distribution of wild *S. bicolor* is narrow compared with *T. triandra* [70], as are the distributions of the 17 native Australian *Sorghum* species, being largely restricted to northern Australia [16]. The chromosomal synteny between these two sentinel *Sorghum* and *Themeda* species provide a route to identify adaptive variation in *T. triandra* and link it to functionally characterised loci in *S. bicolor* (and other well-characterised genera), effectively using *T. triandra* as a discovery platform for climate adaptation genes within the Andropogoneae.

Common-environment (glasshouse) experiments have shown that the *Themeda* accessions from warmer environments take significantly longer to flower, with this effect most pronounced under cool growing conditions, while accessions from drier environments flower earlier, likely as a drought escape strategy [8]. Orthology with well-characterised *S. bicolor* flowering genes [63] provides a way to nominate candidate loci for this variation, although the causal basis remains untested. In the southernmost (Tasmanian) accession of *T. triandra*, genetic differences were concentrated around regulators of the floral repressor *FLC*, along with a mutation in the vernalization-responsive gene *SbVRN1*, a plausible candidate given its known role as the central determinant of the vernalisation requirement for temperate cereals to survive colder, shorter growing seasons [71]. The PAN accession, collected from an environment where spasmodic rain events impose pressure to reproduce opportunistically during transient, favourable conditions, carried high-impact mutations across a much broader set of candidate genes spanning photoperiod response, floral signalling and transition pathways. These contrasting genetic adaptations in the flowering pathway between southern and northern *T. triandra* accessions are consistent with distinct hypotheses for adaptation of flowering time to local environment and climate, although functional validation (e.g. expression experiments, QTL/GWAS, or transgenic work) is required to confirm any causal role. Critically, these candidate genes were identified through direct orthology with *Sorghum*, illustrating how the syntenic relationship between these genomes can translate ecological variation in *T. triandra* into candidate genes for crop improvement, once validated.

Flowering time represents just one of many trait systems that can now be explored using this resource. Although gene family enrichment patterns were too broad to attribute to specific selective pressures, *T. triandra* families absent in *Sorghum/Zea* and variable-copy families were enriched for stress and defence signalling, redox chemistry, secondary metabolism and cell wall modification. Since incomplete annotation would only remove such families from this set (and not add to them) this is likely to understate the lineage-specific gene content. These functions could potentially underpin this species’ success across its diverse climate range. Many of these same pathways are targets for crop improvement in sorghum, where narrowed genetic diversity in stress-responsive gene families has been observed in domesticated material [69, 72]. Of particular interest are traits that may have been selected against during domestication due to trade-offs with yield, but which remain intact in wild species. For example, disease resistance genes have been progressively lost during domestication in multiple crops, with each additional nucleotide-binding leucine-rich repeat (NLR) receptor gene estimated to impose an average 1.6% yield penalty [73, 74]. Similarly, cuticular wax composition affects both drought tolerance and pathogen defence in many plants and has diverged substantially, even within wild species, over short evolutionary timescales [75, 76] and this trait has been further modified during crop improvement [77, 78]. Combined with the syntenic framework described here, the lineage-specific gene content of *T. triandra* provides a starting point for identifying candidate genes for climate resilience that may have been lost or diminished in the cultivated sorghum gene pool.

This information also has direct applications for conservation and management of *T. triandra* itself. Temperate grasslands dominated by *T. triandra* are among Australia’s most endangered ecosystems [79], and restoration efforts are complicated by the species’ low seed viability and limited understanding of local adaptation [80]. The phylogeographic structure described here, combined with the ploidy variation across populations, has practical implications for sourcing seed for restoration. Our results show that diploid populations from cool, wet environments diverge strongly at the genome level from those in the hot arid north-west; matching provenance to restoration site is likely to matter for establishment success [81]. Ploidy variation adds a further layer of complexity, with polyploidy substantially reducing predicted genomic vulnerability to future climates [11]. Although our sampling captures only a small subset of each population, polyploids largely dominated the warmer inland environments, as has been observed previously [11]. However, an interesting observation is that the population of *T. triandra* sampled from the Pilbara region in WA (PAN), the most arid site, was diploid. Previous samples taken from this region were found to be polyploid, with a chloroplast haplotype similar to that of those from the south-east of Australia suggestive of ancient chloroplast capture [10]. This particular population is notable, challenging the expectation that polyploidy is the default evolutionary response to greater environmental stress in *T. triandra* [82].

The reproduction of *T. triandra* combines wind-pollinated outcrossing with self-pollination and apomixis [83]. This has important implications both for natural adaptation and restoration applications. Multiple reproductive mechanisms in *T. triandra* enable it to be used for genetic analysis through controlled crosses and segregation studies, making it possible to link adaptive traits to specific genotypes. The reference genome and SNP framework presented here now make it possible to move beyond neutral markers toward identifying the specific genes and variants associated with climate adaptation in different *T. triandra* populations, enabling more informed decisions about seed sourcing, assisted migration, and genetic rescue as conditions shift under climate change.

The chromosome-scale reference genome presented here, together with resequencing from climatic extremes and broad phylogeographic sampling from both this study and others [10, 11], establishes *T. triandra* as a genomic resource for investigating climate adaptation in a wild Andropogoneae grass. This work opens several directions for further investigation. Resequencing additional accessions, particularly polyploid populations and those from regions with unique soil and microclimatic profiles, would test whether the patterns of adaptive variation hold across the species broader range; investing in long reads for these analysis would allow for stronger inference of copy number and structural variation than short reads can provide (due to divergence and heterozygosity impacting mapping efficiency and obfuscating true copy loss). A pan-genome incorporating multiple *Themeda* accessions (and species) would capture presence-absence variation not accessible from a single reference, and combining this with existing *Sorghum* and *Zea* pangenomes could establish a tribe-level comparative resource for the Andropogoneae [84], which would offer vast insights into adaptive diversity, given the over 1200 species spanning ∼20 million years of divergence [85]. The resources presented here provide both a foundation for conservation genomics and a rich source of candidate genes shaped by millions of years of selection across some of the most contrasting environments occupied by any single grass species.

## ACKNOWLEDGEMENTS

This was a joint project of the Australian Research Council Centre of Excellence for Plant Success in Nature and Agriculture (CE200100015) and Bioplatforms Australia. We acknowledge the contribution of the Australian Grasslands Initiative Consortium in the generation of data used in this publication. The Initiative is supported by funding from Bioplatforms Australia, enabled by the Commonwealth Government National Collaborative Research Infrastructure Strategy (NCRIS). We also acknowledge the provision of computing and data resources provided by the Australian BioCommons Leadership Share (ABLeS) program, which is co-funded by Bioplatforms Australia (enabled by NCRIS), the National Computational Infrastructure and Pawsey Supercomputing Research Centre. We also acknowledge the use of the high-performance computing facilities provided by Digital Research Services, IT Services at the University of Tasmania. We thank Juanita Christine Lauer, Marne Marie Dunin and Rachel Burton (University of Adelaide), Caroline Cristofolini (Royal Botanic Gardens Sydney) and Daniel Pascoe (Macquarie University) for assistance with cell cytometry measurements. The reference plant (WBW) was collected by Rachael Gallagher (Western Sydney University). Useful comments on ploidy data by Richard Frankham (Macquarie University) are appreciated.

## AUTHOR CONTRIBUTIONS

BJA coordinated funding resources for sequencing, conceptualized the project and assembled the germplasm collection with VKJ, JLH, ACL and EF. VKJ, JLH and BJA collected experimental material for ACL and LC to complete extractions and sample preparation. VKJ, JLH and BJA managed sample dispatch to analytical service laboratories via Bioplatforms facilities. TRA and JBB performed the bioinformatics analysis on all sequencing data and BJA provided the cytometry data in discussion with EF, ACL and JLJ. The manuscript was written by JLH, SMS, JBB and BJA with editing by IJW, EF and VKJ and commentary on a late draft by JLJ and LC.

## DATA STATEMENT

Scripts used to assemble the *Themeda* genome are available at https://github.com/Royal-Botanic-Gardens-Victoria/Themeda. The genome assembly and sequencing data (including ddRAD) are available at the NCBI Sequence Read Archive under BioProject accession PRJNA1433027, with additional annotation files and results available at Zenodo (doi: 10.5281/zenodo.23029265).

## SUPPORTING INFORMATION

**Supplementary Table 1:** *Themeda triandra* accessions sampled for this research

**Supplementary Table 2:** Genome assembly statistics for the *Themeda triandra* WBW v1.0 reference genome

**Supplementary Table 3**: Repeat statistics for the *Themeda triandra* WBW v1.0 reference genome

**Supplementary Table 4:** Comparison of the *Themeda triandra* WBW v1.0 reference assembly with the two previously published *T. triandra* assemblies

**Supplementary Table 5**: Enrichment of GO terms associated with *Themeda triandra* gene families that are unique (a), expanded (b) and contracted (c) compared to *Zea mays* and *Sorghum bicolor*

**Supplementary Table 6:** Enrichment of GO terms associated with *Themeda triandra* genes with high-impact mutations and copy number variation between SBC, SYD and PAN accessions

**Supplementary Figure 1:** Whole-genome alignment of the two previously published *Themeda triandra* assemblies to the WBW v1.0 reference

**Supplementary Figure 2:** Distribution of putative structural variants across *Themeda triandra* chromosomes, based on resequencing of six accessions

**Supplementary Figure 3:** Synteny of the *Themeda triandra* WBW v1.0 genome with *Sorghum bicolor* (BTx623 v3.0)

**Supplementary Figure 4:** Distribution of heterozygous allele read depths across *Themeda triandra* accessions

**Supplementary Figure 5:** Flow cytometry traces for representative diploid, tetraploid and hexaploid *Themeda triandra* accessions

**Supplementary Figure 6:** GO terms enrichment in variable gene sets identified through comparison of diploid *Themeda triandra* accessions

## REFERENCES

1. Arthan, W., et al. Phylogenomics of Andropogoneae (Panicoideae: Poaceae) of Mainland Southeast Asia. Syst. Bot. 42, 418–431 (2017)

2. Holder, F., A phylogenetic study of the South African representatives of the tribe Andropogoneae (Poaceae). 2003, University of the Free State.

3. Dunning, L.T., et al. The recent and rapid spread of Themeda triandra. Botany Letters 164, 327–337 (2017)

4. Arthan, W., et al. Complex evolutionary history of two ecologically significant grass genera, Themeda and Heteropogon (Poaceae: Panicoideae: Andropogoneae). Bot. J. Linn. Soc. 196, 437–455 (2021)

5. Hayman, D. The distribution and cytology of the chromosome races of *Themeda australis* in southern Australia. Aust. J. Bot. 8, 58–68 (1960)

6. Groves, R. Growth and development of five populations of *Themeda australis* in response to temperature. Aust. J. Bot. 23, 951–963 (1975)

7. Evans, L. and R. Knox. Environmental control of reproduction in *Themeda australis*. Aust. J. Bot. 17, 375–389 (1969)

8. Jacob, V., et al. Trait–climate relations in *Themeda triandra*: a widely distributed C_4_ grass and crop wild relative. bioRxiv (2026)

9. Cavanagh, A.M., R.C. Godfree, and J.W. Morgan. Awn length variation in Australia’s most widespread grass, *Themeda triandra*, across its distribution. Aust. J. Bot. 72 (2024)

10. Dunning, L.T., et al. Hybridisation and chloroplast capture between distinct Themeda triandra lineages in Australia. Mol. Ecol. 31, 5846–5860 (2022)

11. Ahrens, C.W., et al. Spatial, climate and ploidy factors drive genomic diversity and resilience in the widespread grass *Themeda triandra*. Mol. Ecol. 29, 3872–3888 (2020)

12. Estep, M.C., et al. Allopolyploidy, diversification, and the Miocene grassland expansion. PNAS 111, 15149–15154 (2014)

13. Birari, S.P. Polyploidy in species of *Themeda* Forsk. Caryologia 34, 301–310 (1981)

14. Godfree, R.C., et al. Empirical evidence of fixed and homeostatic patterns of polyploid advantage in a keystone grass exposed to drought and heat stress. Royal Society Open Science 4 (2017)

15. Stevens, A.V., et al. Polyploidy affects the seed, dormancy and seedling characteristics of a perennial grass, conferring an advantage in stressful climates. Plant Biol. 22, 500–513 (2020)

16. Ananda, G.K.S., et al. Wild sorghum as a promising resource for crop improvement. Front Plant Sci. Volume 11 -2020 (2020)

17. Zhu, Y., et al. HEMU: An integrated comparative genomics database and analysis platform for Andropogoneae grasses. Plant Communications 5, 100786 (2024)

18. Karger, D.N., et al. Climatologies at high resolution for the earth’s land surface areas. Sci. Data 4, 170122 (2017)

19. Cheng, H., et al. Haplotype-resolved de novo assembly using phased assembly graphs with hifiasm. Nat. Methods 18, 170–175 (2021)

20. Roach, M.J., S.A. Schmidt, and A.R. Borneman. Purge Haplotigs: allelic contig reassignment for third-gen diploid genome assemblies. BMC Bioinformatics 19, 460 (2018)

21. Marçais, G. and C. Kingsford. A fast, lock-free approach for efficient parallel counting of occurrences of k-mers. Bioinformatics 27, 764–770 (2011)

22. Ranallo-Benavidez, T.R., K.S. Jaron, and M.C. Schatz. GenomeScope 2.0 and Smudgeplot for reference-free profiling of polyploid genomes. Nat. Commun. 11, 1432 (2020)

23. Zhou, C., S.A. McCarthy, and R. Durbin. YaHS: yet another Hi-C scaffolding tool. Bioinformatics 39 (2022)

24. Yates, A.D., et al. Ensembl 2026. Nucleic Acids Res. 54, D1053–D1060 (2026)

25. Astashyn, A., et al. Rapid and sensitive detection of genome contamination at scale with FCS-GX. Genome Biol. 25, 60 (2024)

26. Simão, F.A., et al. BUSCO: assessing genome assembly and annotation completeness with single-copy orthologs. Bioinformatics 31, 3210–3212 (2015)

27. Huang, N. and H. Li. compleasm: a faster and more accurate reimplementation of BUSCO. Bioinformatics 39 (2023)

28. Brown, M.R., P. Manuel Gonzalez de La Rosa, and M. Blaxter. tidk: a toolkit to rapidly identify telomeric repeats from genomic datasets. Bioinformatics 41 (2025)

29. Su, W., et al., A tutorial of EDTA: extensive De novo TE annotator, in Plant Transposable Elements: Methods and Protocols. 2021, Humana: New York. p. 55–67.

30. Gabriel, L., et al. BRAKER3: Fully automated genome annotation using RNA-seq and protein evidence with GeneMark-ETP, AUGUSTUS, and TSEBRA. Genome Res. 34, 769–777 (2024)

31. Paysan-Lafosse, T., et al. InterPro in 2022. Nucleic Acids Res. 51, D418–D427 (2023)

32. Cantalapiedra, C.P., et al. eggNOG-mapper v2: Functional annotation, orthology assignments, and domain prediction at the metagenomic scale. Mol. Biol. Evol. 38, 5825–5829 (2021)

33. Kanehisa, M., Y. Sato, and K. Morishima. BlastKOALA and GhostKOALA: KEGG tools for functional characterization of genome and metagenome sequences. J. Mol. Biol. 428, 726–731 (2016)

34. Li, H. Minimap2: pairwise alignment for nucleotide sequences. Bioinformatics 34, 3094–3100 (2018)

35. McCormick, R.F., et al. The Sorghum bicolor reference genome: improved assembly, gene annotations, a transcriptome atlas, and signatures of genome organization. Plant J 93, 338–354 (2018)

36. Marçais, G., et al. MUMmer4: A fast and versatile genome alignment system. PLoS Comp. Biol. 14, e1005944 (2018)

37. Goel, M., et al. SyRI: finding genomic rearrangements and local sequence differences from whole-genome assemblies. Genome Biol. 20, 1–13 (2019)

38. Sun, J., et al. OrthoVenn3: an integrated platform for exploring and visualizing orthologous data across genomes. Nucleic Acids Res. 51, W397–W403 (2023)

39. Schnable, P.S., et al. The B73 maize genome: complexity, diversity, and dynamics. Science 326, 1112–1115 (2009)

40. Li, L., C.J. Stoeckert, Jr., and D.S. Roos. OrthoMCL: identification of ortholog groups for eukaryotic genomes. Genome Res. 13, 2178–89 (2003)

41. Mendes, F.K., et al. CAFE 5 models variation in evolutionary rates among gene families. Bioinformatics 36, 5516–5518 (2020)

42. Wang, Y., et al. MCScanX: a toolkit for detection and evolutionary analysis of gene synteny and collinearity. Nucleic Acids Res. 40, e49–e49 (2012)

43. Katoh, K. and D.M. Standley. MAFFT multiple sequence alignment software version 7: Improvements in performance and usability. Mol. Biol. Evol. 30, 772–80 (2013)

44. Suyama, M., D. Torrents, and P. Bork. PAL2NAL: robust conversion of protein sequence alignments into the corresponding codon alignments. Nucleic Acids Res. 34, W609–W612 (2006)

45. Yang, Z. PAML 4: Phylogenetic Analysis by Maximum Likelihood. Mol. Biol. Evol. 24, 1586–1591 (2007)

46. Doyle, J. and J. Doyle. A rapid DNA isolation procedure for small quantities of fresh leaf tissue. Phytochem. Bull. 19, 11–15 (1987)

47. Chen, S., et al. fastp: an ultra-fast all-in-one FASTQ preprocessor. Bioinformatics 34, i884–i890 (2018)

48. Catchen, J., et al. Stacks: an analysis tool set for population genomics. Mol. Ecol. 22, 3124–3140 (2013)

49. Krueger, F., S.R. Andrews, and C.S. Osborne. Large scale loss of data in low-diversity Illumina sequencing libraries can be recovered by deferred cluster calling. PLoS One 6, e16607 (2011)

50. Bolger, A.M., M. Lohse, and B. Usadel. Trimmomatic: a flexible trimmer for Illumina sequence data. Bioinformatics 30, 2114–2120 (2014)

51. Li, H. Aligning sequence reads, clone sequences and assembly contigs with BWA-MEM. arXiv 1303.3997 (2013)

52. Danecek, P., et al. Twelve years of SAMtools and BCFtools. Gigascience 10, giab008 (2021)

53. Chen, X., et al. Manta: rapid detection of structural variants and indels for germline and cancer sequencing applications. Bioinformatics 32, 1220–1222 (2015)

54. Weißbach, S., et al. Reliability of genomic variants across different next-generation sequencing platforms and bioinformatic processing pipelines. BMC Genom. 22, 62 (2021)

55. Severn-Ellis, A.A., et al., Genotyping for species identification and diversity assessment using double-digest restriction site-associated DNA sequencing (ddRAD-Seq), in Legume genomics: Methods and protocols. 2020, Springer US: New York. p. 159–187.

56. Garrison, E. and G. Marth. Haplotype-based variant detection from short-read sequencing. arXiv 1207.3907 (2012)

57. Kozlov, A.M., et al. RAxML-NG: a fast, scalable and user-friendly tool for maximum likelihood phylogenetic inference. Bioinformatics 35, 4453–4455 (2019)

58. Revell, L.J. phytools 2.0: an updated R ecosystem for phylogenetic comparative methods (and other things). PeerJ 12, e16505 (2024)

59. Talevich, E., et al. CNVkit: Genome-wide copy number detection and visualization from targeted DNA sequencing. PLoS Comp. Biol. 12, e1004873 (2016)

60. Cingolani, P., et al. A program for annotating and predicting the effects of single nucleotide polymorphisms, SnpEff. Fly 6, 80–92 (2012)

61. Alexa, A. and J. Rahnenfuhrer. topGO: Enrichment analysis for gene ontology. R package version 2.64.0 https://bioconductor.org/packages/topGO (2023)

62. Swigoňová, Z., et al. Close split of sorghum and maize genome progenitors. Genome Res. 14, 1916–1923 (2004)

63. Mace, E.S., C.H. Hunt, and D.R. Jordan. Supermodels: sorghum and maize provide mutual insight into the genetics of flowering time. Theor. Appl. Genet. 126, 1377–1395 (2013)

64. Michaels, S.D. and R.M. Amasino. FLOWERING LOCUS C encodes a novel MADS domain protein that acts as a repressor of flowering. Plant Cell 11, 949–956 (1999)

65. Sheldon, C.C., et al. The molecular basis of vernalization: The central role of *FLOWERING LOCUS C* (*FLC*). PNAS 97, 3753–3758 (2000)

66. Burgarella, C., et al. The road to sorghum domestication: Evidence from nucleotide diversity and gene expression patterns. Front Plant Sci. Volume 12–2021 (2021)

67. Casa, A.M., et al. Diversity and selection in sorghum: simultaneous analyses using simple sequence repeats. Theor. Appl. Genet. 111, 23–30 (2005)

68. Mace, E.S., et al. Whole-genome sequencing reveals untapped genetic potential in Africa’s indigenous cereal crop sorghum. Nat. Commun. 4, 2320 (2013)

69. Tack, J., J. Lingenfelser, and S.V.K. Jagadish. Disaggregating sorghum yield reductions under warming scenarios exposes narrow genetic diversity in US breeding programs. PNAS 114, 9296–9301 (2017)

70. Mace, E.S., et al. A global resource for exploring and exploiting genetic variation in sorghum crop wild relatives. Crop Sci. 61, 150–162 (2021)

71. Yan, L., et al. Positional cloning of the wheat vernalization gene *VRN1*. PNAS 100, 6263–6268 (2003)

72. Massel, K., et al. Whole genome sequencing reveals potential new targets for improving nitrogen uptake and utilization in *Sorghum bicolor*. Front Plant Sci. Volume 7 -2016 (2016)

73. Derbyshire, M.C., et al. The complex relationship between disease resistance and yield in crops. Plant Biotechnol. J. 22, 2612–2623 (2024)

74. Gao, M., et al. Revisiting growth–defence trade-offs and breeding strategies in crops. Plant Biotechnol. J. 22, 1198–1205 (2024)

75. Asadyar, L., et al. Evidence for within-species transition between drought response strategies in *Nicotiana benthamiana*. New Phytol. 244, 464–476 (2024)

76. Lewandowska, M., A. Keyl, and I. Feussner. Wax biosynthesis in response to danger: its regulation upon abiotic and biotic stress. New Phytol. 227, 698–713 (2020)

77. Jordan, W.R., et al. Environmental physiology of Sorghum. II. Epicuticular wax load and cuticular transpiration. Crop Sci. 24, cropsci1984.0011183X002400060038x (1984)

78. Xue, D., et al. Molecular and evolutionary mechanisms of cuticular wax for plant drought tolerance. Front Plant Sci. Volume 8 - 2017 (2017)

79. Cole, B.I. and I.D. Lunt. Restoring kangaroo grass (*Themeda triandra*) to grassland and woodland understoreys: a review of establishment requirements and restoration exercises in south-east Australia. Ecol. Manage. Restor. 6, 28–33 (2005)

80. Male, D., et al. Time of seed harvest and sowing determines successful establishment of kangaroo grass (*Themeda triandra*) on Dja Dja Wurrung Country. Crop & Pasture Science 76 (2025)

81. Prober, S.M., et al. Climate-adjusted provenancing: a strategy for climate-resilient ecological restoration. Front. Ecol. Evol. Volume 3 - 2015 (2015)

82. Tossi, V.E., et al. Impact of polyploidy on plant tolerance to abiotic and biotic stresses. Front Plant Sci. Volume 13 - 2022 (2022)

83. Snyman, H.A., L.J. Ingram, and K.P. Kirkman. *Themeda triandra*: a keystone grass species. African Journal of Range & Forage Science 30, 99–125 (2013)

84. Tao, Y., et al. Extensive variation within the pan-genome of cultivated and wild sorghum. Nat. Plants 7, 766–773 (2021)

85. Welker, C.A.D., et al. Phylogenomics enables biogeographic analysis and a new subtribal classification of Andropogoneae (Poaceae—Panicoideae). Journal of Systematics and Evolution 58, 1003–1030 (2020)

