## Supplementary Figures 1-6 and Supplementary Tables 2-3 for "Genome assembly of the C_4_ grass *Themeda triandra* confirms karyotype orthology with sorghum and reveals genetic variation across diverse environments"

**Supplementary Material**

**Supplementary Table 1:** *Themeda triandra* accessions sampled for this research (excel file)

**Supplementary Table 2**: Genome assembly statistics for the *Themeda triandra* WBW v1.0 reference genome

| **Measure** | ***Themeda triandra* WBW v1.0** |
| --- | --- |
| Assembly size | 754,731,078 bp |
| Contig/scaffold count | 25 / 22 |
| Contig/scaffold N50 | 61.719 / 76.500 Mbp |
| GC content | 46.15% |
| Ns  Telomeres | <0.001%  15 |
| BUSCOs (genome, poales_odb10) |  |
| - Single copy | 94.98% (4650) |
| - Duplicated | 4.76% (233) |
| - Fragmented | 0.06% (3) |
| - Missing | 0.20% (10) |
| BUSCOs (annotation, poales_odb10) |  |
| - Single copy | 62.77% (3073) |
| - Duplicated | 11.79% (577) |
| - Fragmented | 1.29% (63) |
| - Missing | 24.16% (1183) |

**Supplementary Table 3**: Repeat statistics for the *Themeda triandra* WBW v1.0 reference genome. Values for LTR retrotransposon subclasses (Copia, Gypsy, unclassified) and DNA transposon superfamilies are subtotals of the class totals above them (marked with *)

| **Repeat class** | **Length (bp)** | **Number of elements** | **Percentage of sequence** |
| --- | --- | --- | --- |
| **Class I retrotransposons (LTR)** | | | |
| LTR retrotransposons | 307,026,830 | 225,849 | 40.68 |
| *(Copia) | 38,543,725 | 26,129 | 5.11 |
| *(Gypsy) | 130,556,038 | 72,943 | 17.30 |
| *(unclassified LTR) | 137,927,067 | 126,777 | 18.28 |
| **Class II DNA transposons** | | | |
| TIR DNA transposons | 53,632,455 | 156,690 | 7.11 |
| *(CACTA) | 18,560,003 | 43,020 | 2.46 |
| *(Mutator) | 9,676,217 | 26,380 | 1.28 |
| *(PIF/Harbinger) | 14,484,654 | 46,812 | 1.92 |
| *(Tc1/Mariner) | 7,794,583 | 30,498 | 1.03 |
| *(hAT) | 3,116,998 | 9,980 | 0.41 |
| Helitrons/Rolling circle | 76,487,146 | 162,606 | 10.13 |
| **Other elements** | | | |
| Unclassified repeat fragments | 1,447,457 | 7,095 | 0.19 |
| **Total interspersed repeats** | 438,593,888 | 552,240 | 58.11 |

**Supplementary Table 4:** Comparison of the *Themeda triandra* WBW v1.0 reference assembly with the two previously published *T. triandra* assemblies (excel file)

**Supplementary Table 5**: Enrichment of GO terms associated with *Themeda triandra* gene families that are unique (a), expanded (b) and contracted (c) compared to *Zea mays* and *Sorghum bicolor* (excel file)

**Supplementary Table 6**: Enrichment of GO terms associated with *Themeda triandra* genes with high-impact mutations and copy number variation between SBC, SYD and PAN accessions (excel file)

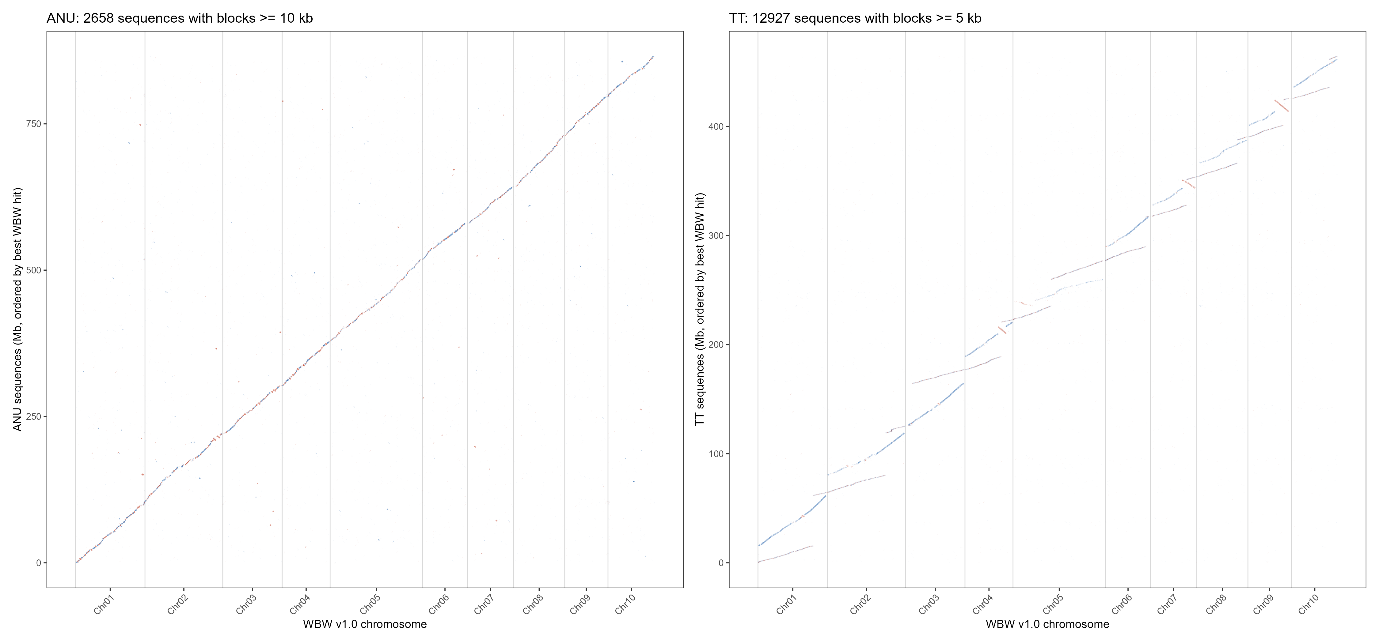

**Supplementary Figure 1:** Whole-genome alignment of the two previously published *Themeda triandra* assemblies to the WBW v1.0 reference. Alignments of (a) GCA_018135685.1 and (b) GCA_025603265.1 against the ten WBW pseudomolecules (minimap2 -x asm20; primary alignments with MAPQ ≥ 5 and blocks ≥ 10 kb in (a), ≥ 5 kb in (b), for plotting). Blue and red denote same- and opposite-strand alignments, but the orientation of individual contigs is arbitrary.

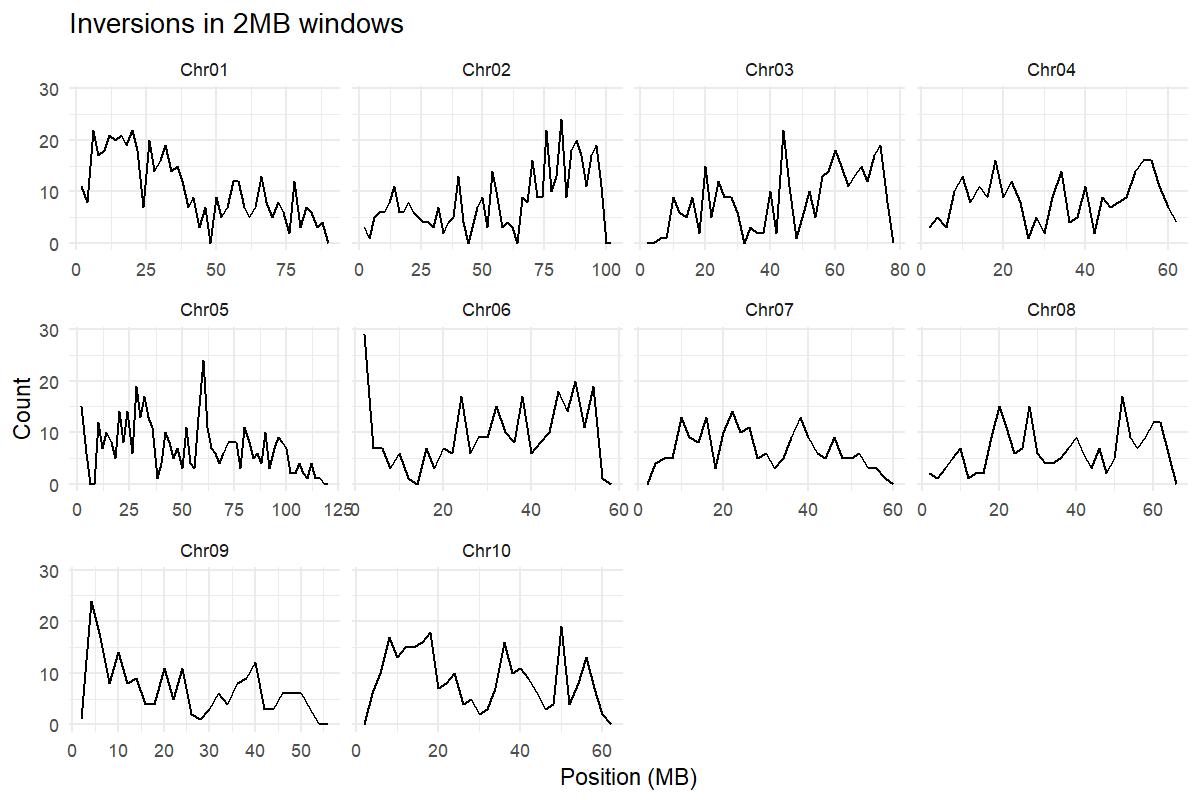

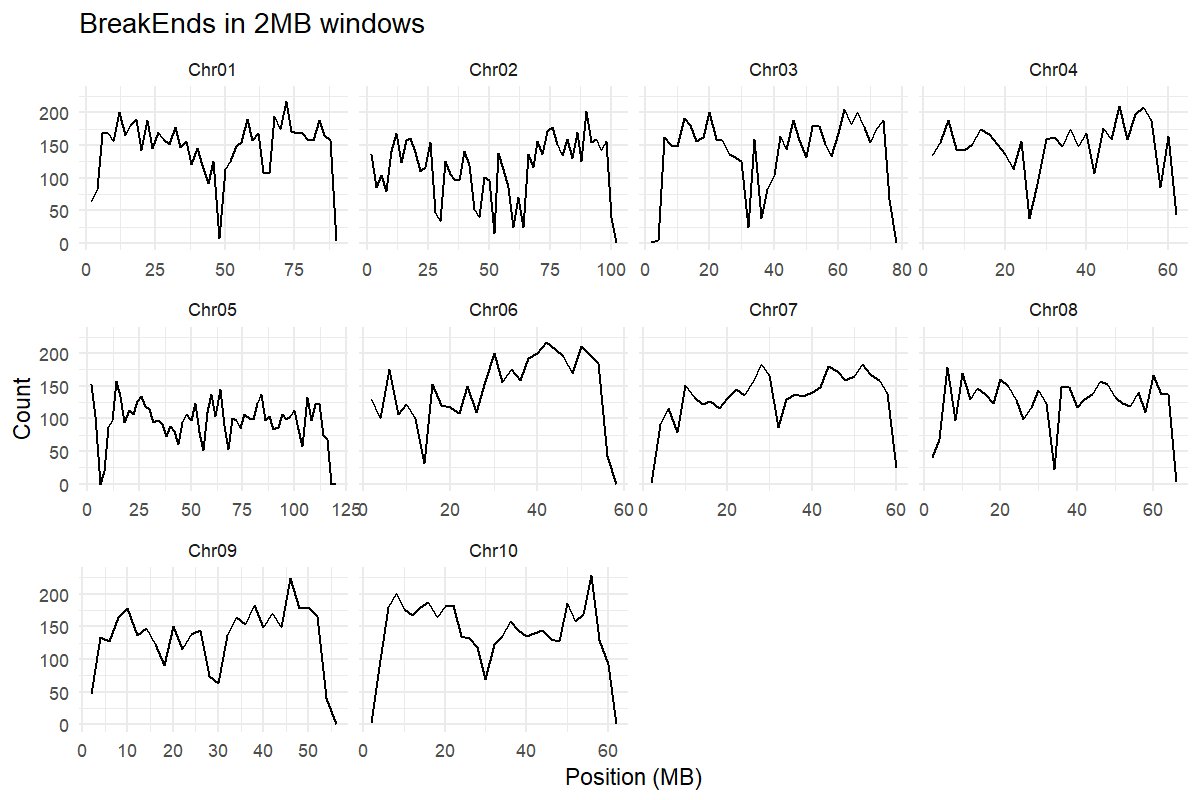

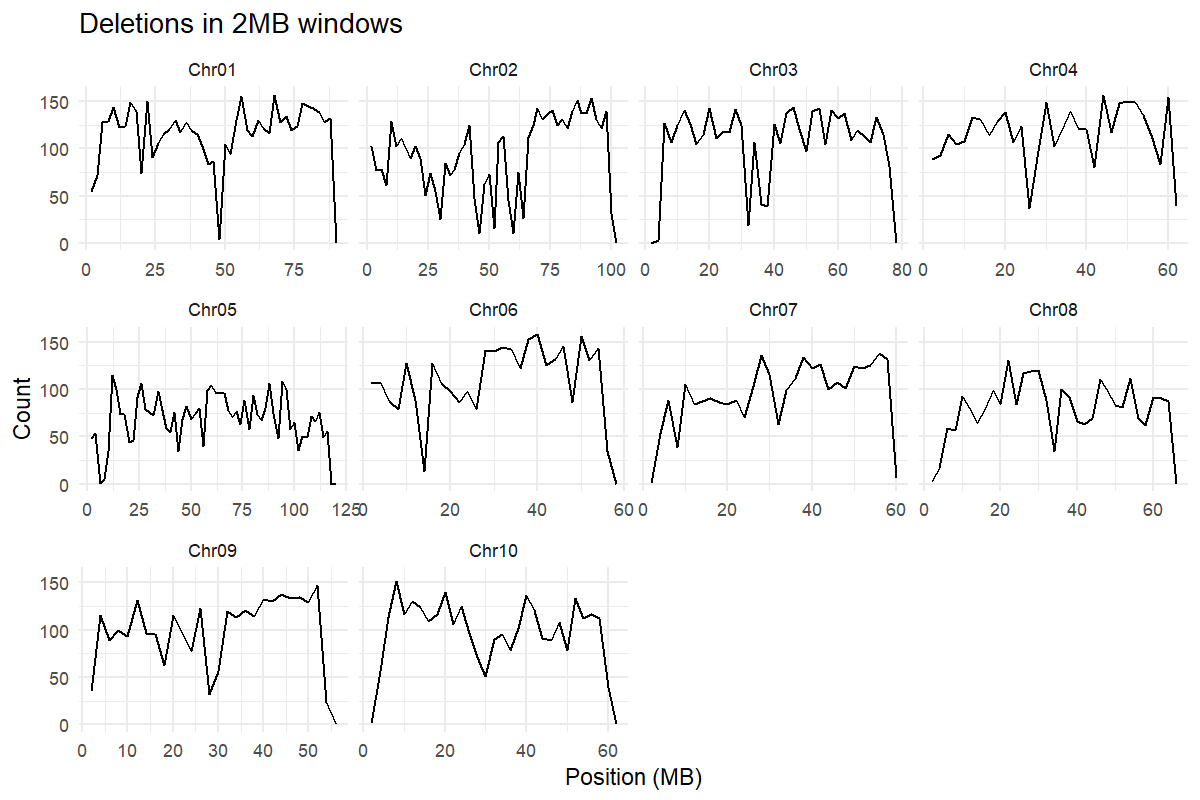

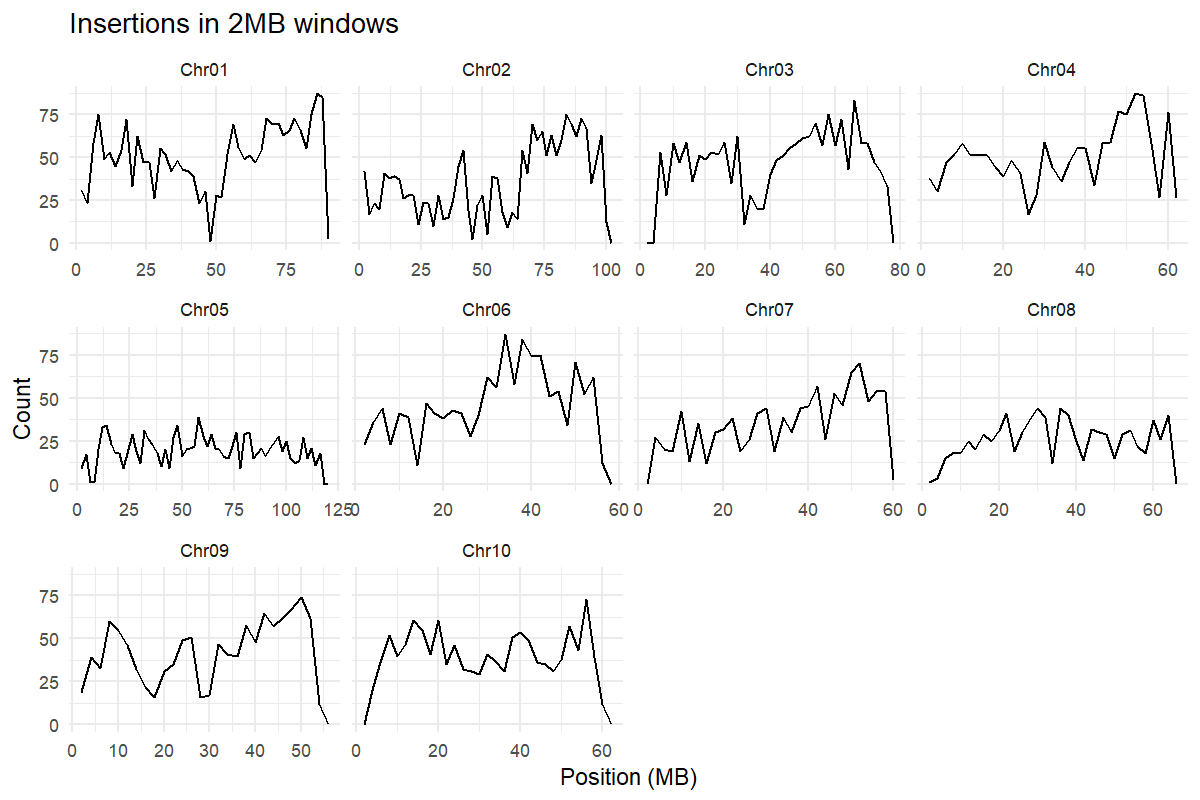

**Supplementary Figure 2**: Distribution of putative structural variants across *Themeda* *triandra* chromosomes, based on resequencing of six accessions. Summary of *manta* output, grouping putative structural variants into counts within 2 Mbp windows.

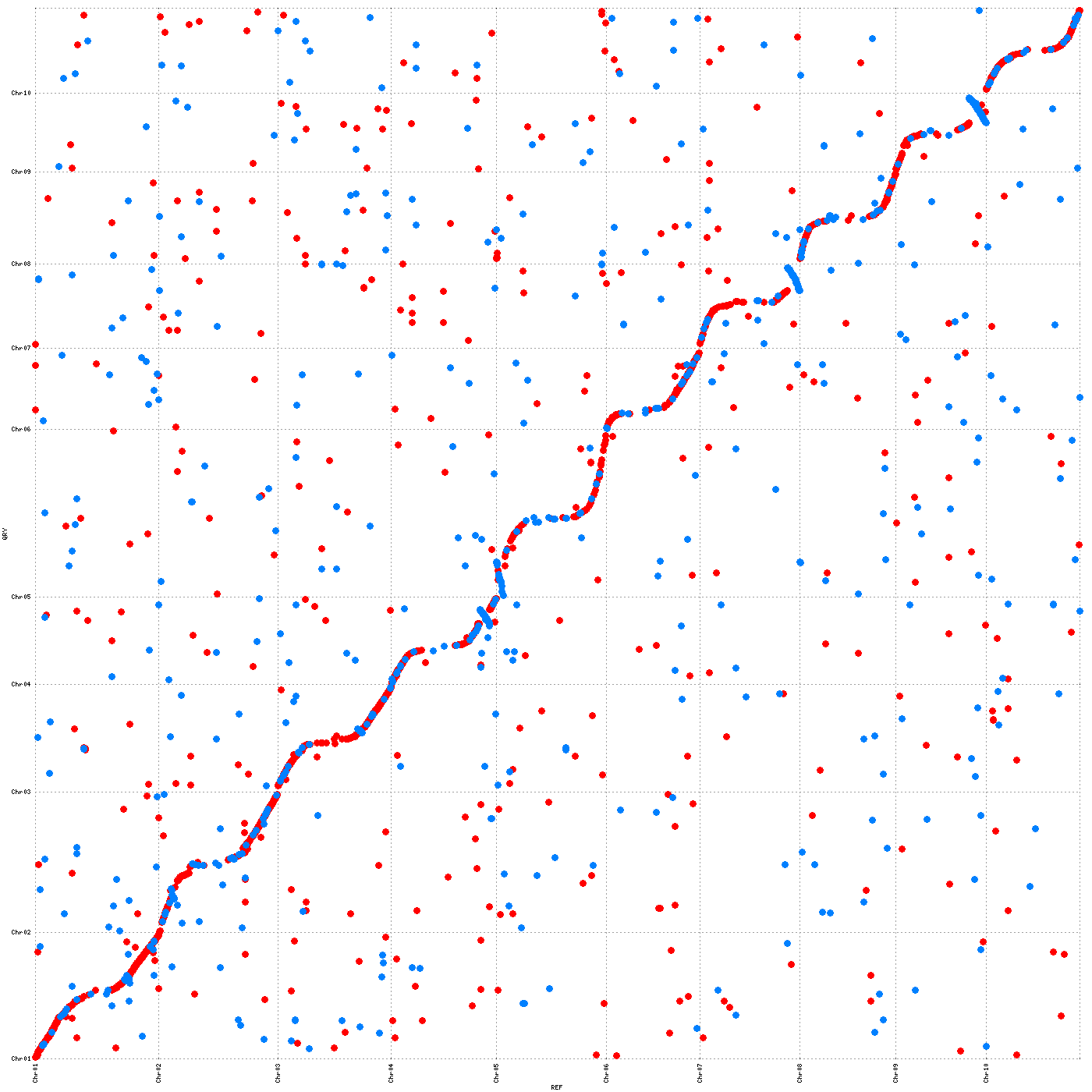

**Supplementary Figure 3:** Synteny of the *Themeda triandra* WBW v1.0 genome with *Sorghum bicolor* (BTx623 v3.0). Points indicate segments of DNA sequence between 103 – 13,015 bp (average 2,719 bp) that were found in both genome assemblies, with blue and red denoting same- and opposite-strand alignments.

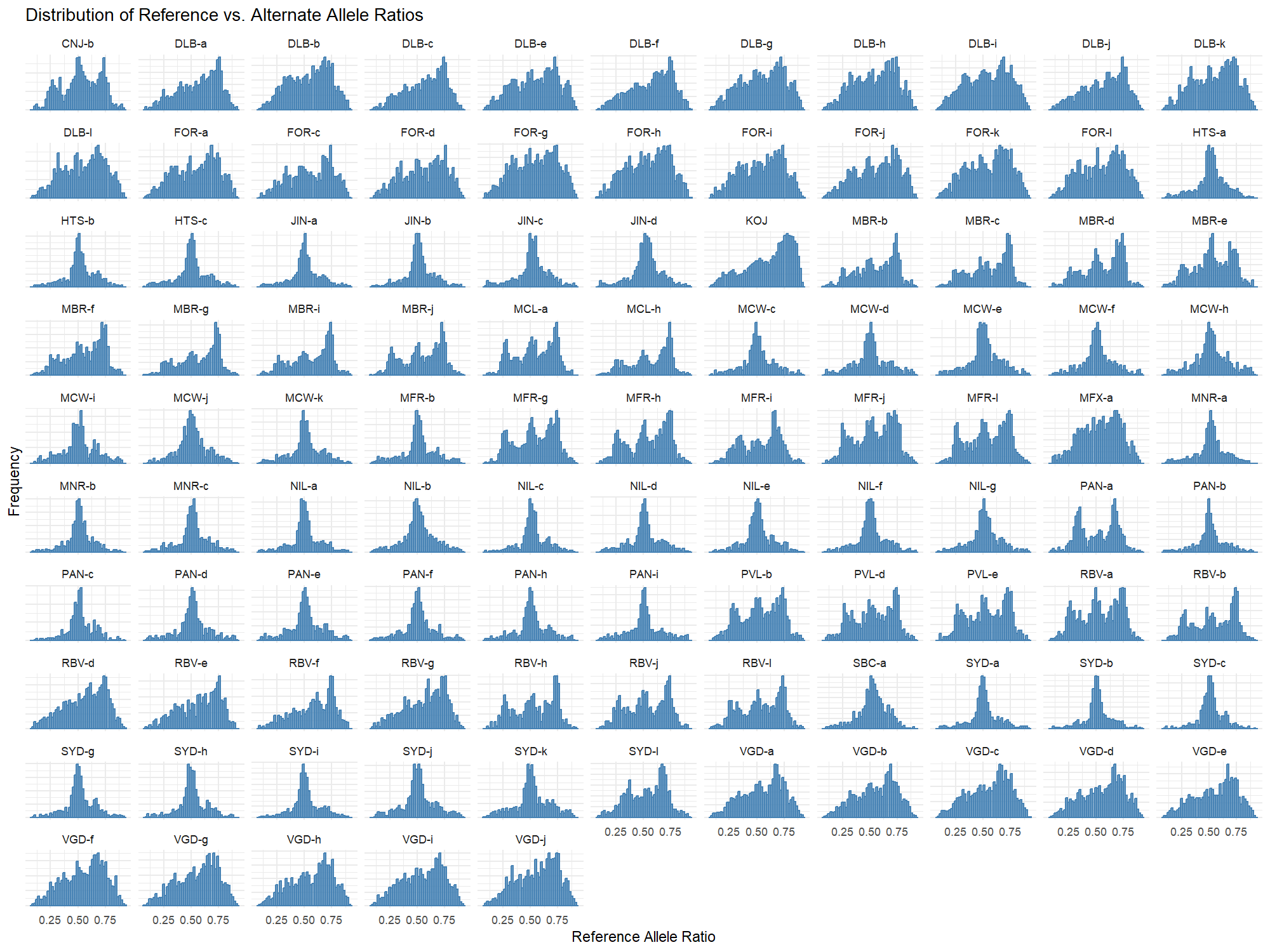

**Supplementary Figure 4:** Distribution of heterozygous allele read depths across *Themeda triandra* accessions. Putative ploidy is indicated by number of peaks +1. Right skewed graphs are putative hexaploids based on association with cytometry results.

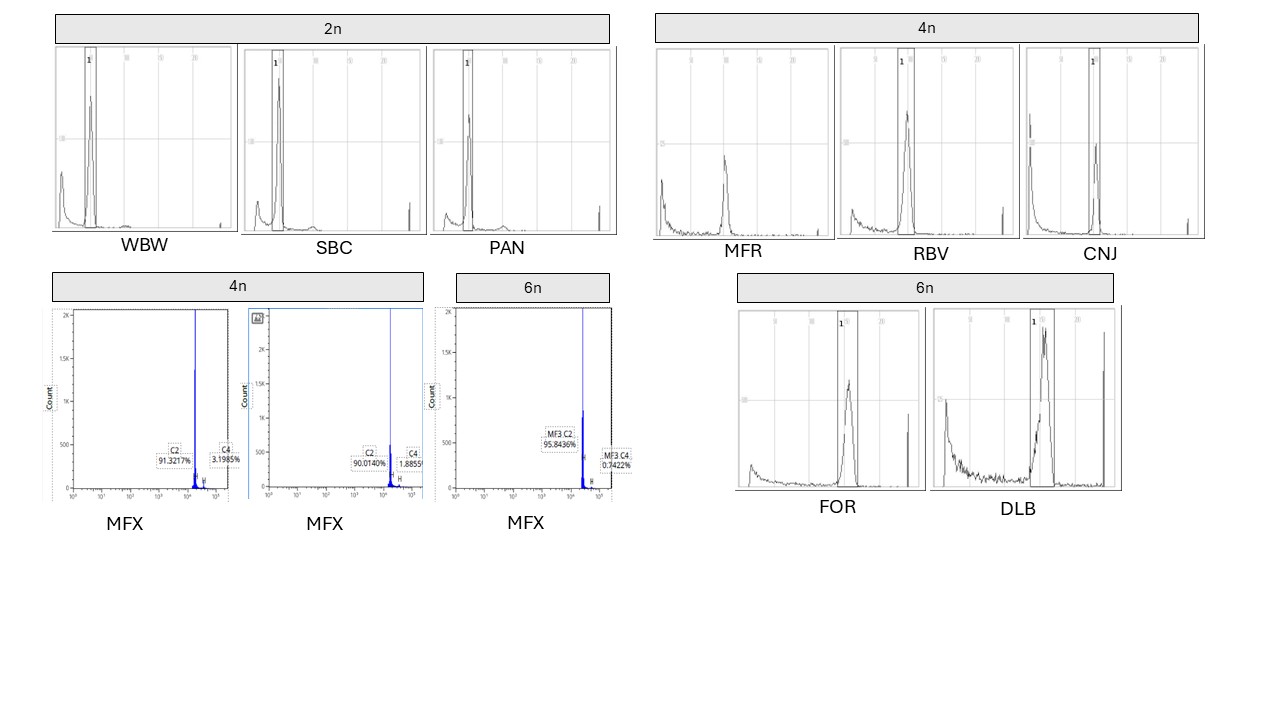

**Supplementary Figure 5:** Flow cytometry traces for representative diploid, tetraploid and hexaploid *Themeda triandra* accessions (see complete ploidy analysis in Figure 4 and Supplementary Figure 4). Reference accessions were used to establish ploidy peaks, namely a diploid accession from Tasmania and a tetraploid accession from Winderlup, South Australia (see Methods). The reference accession (‘WBW’) is shown to be diploid. Flow cytometry was performed in two laboratories, with three Mt Fox (55830) accessions revealing both diploid and tetraploid individuals.

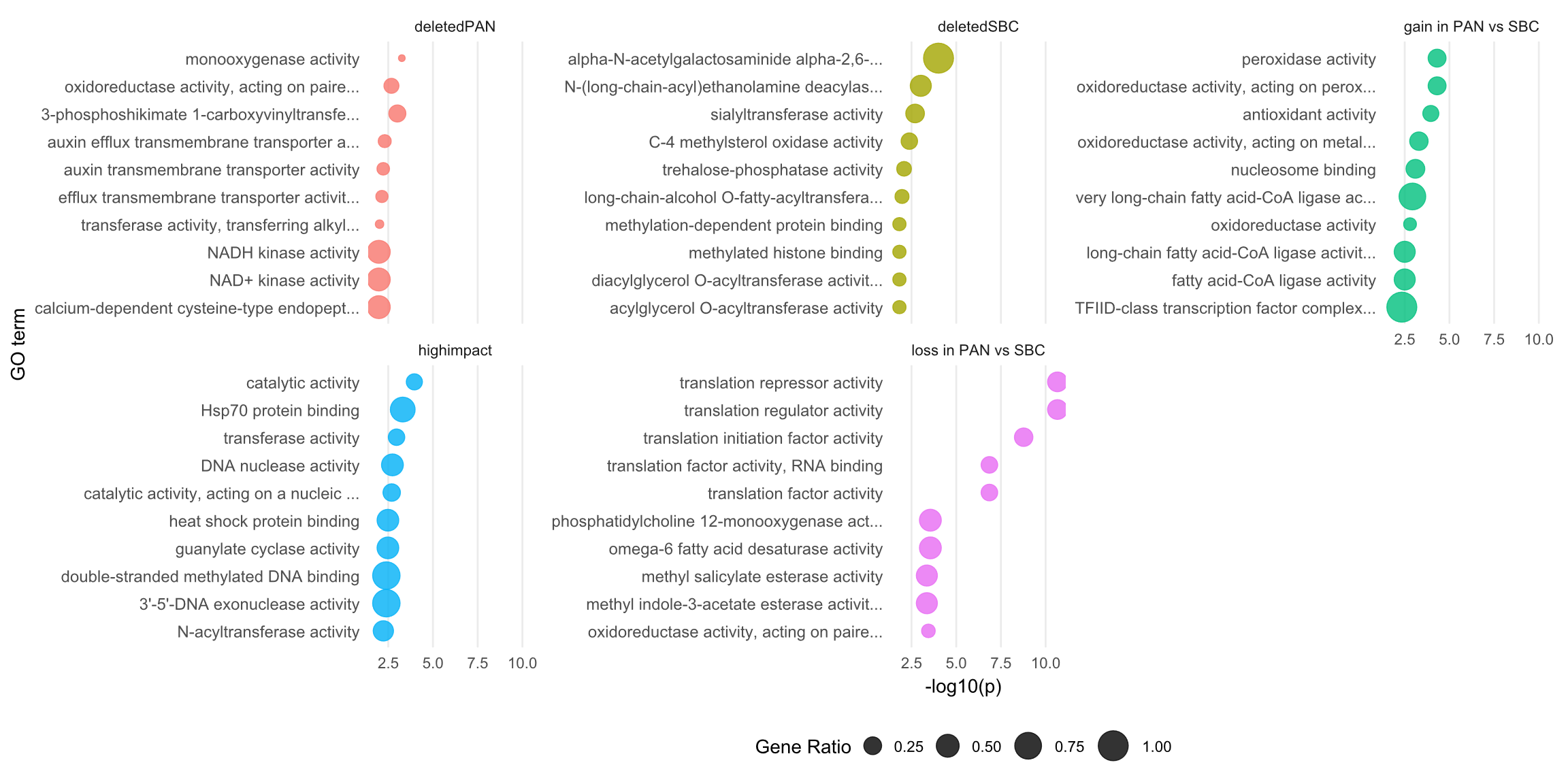

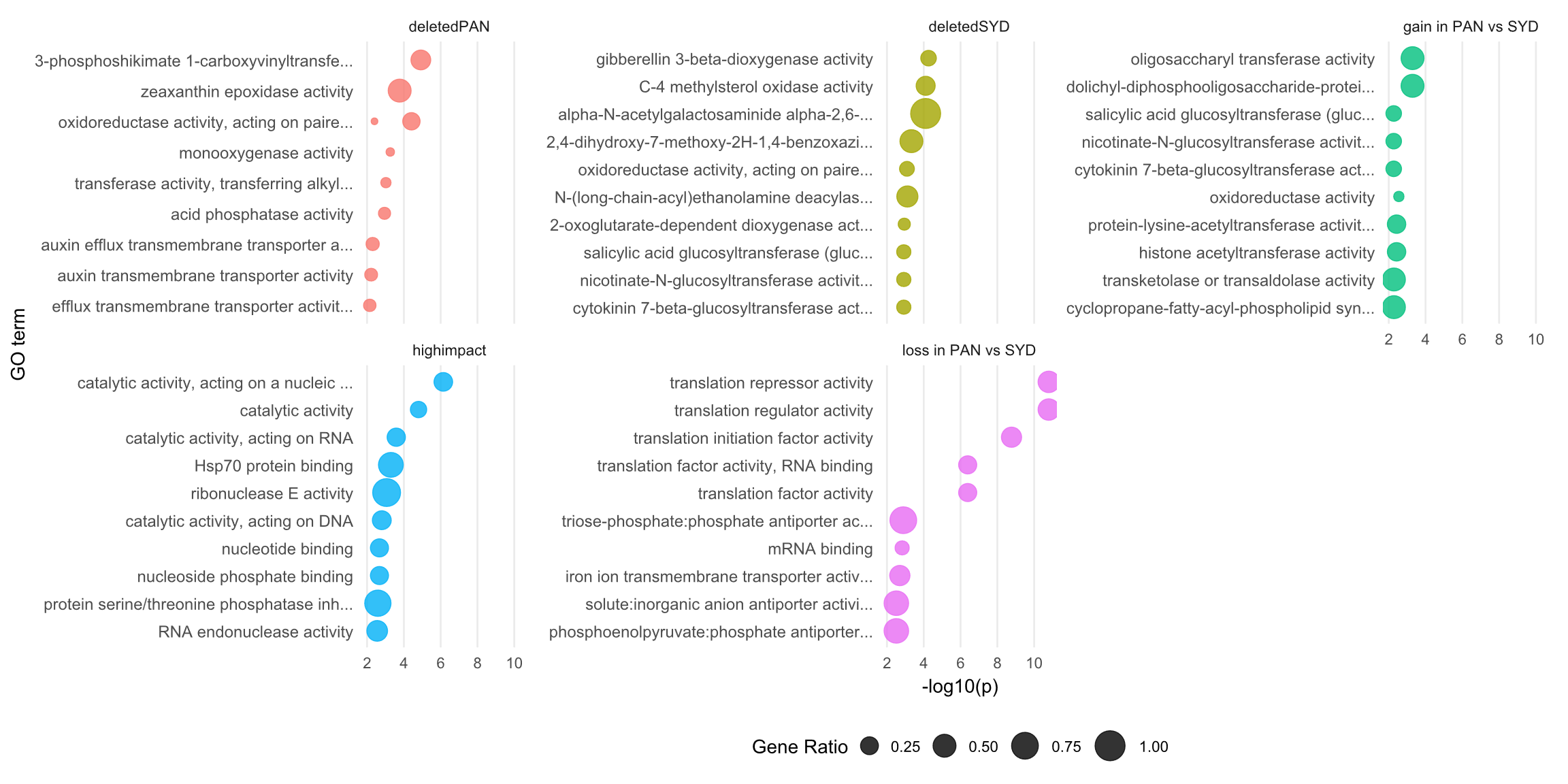

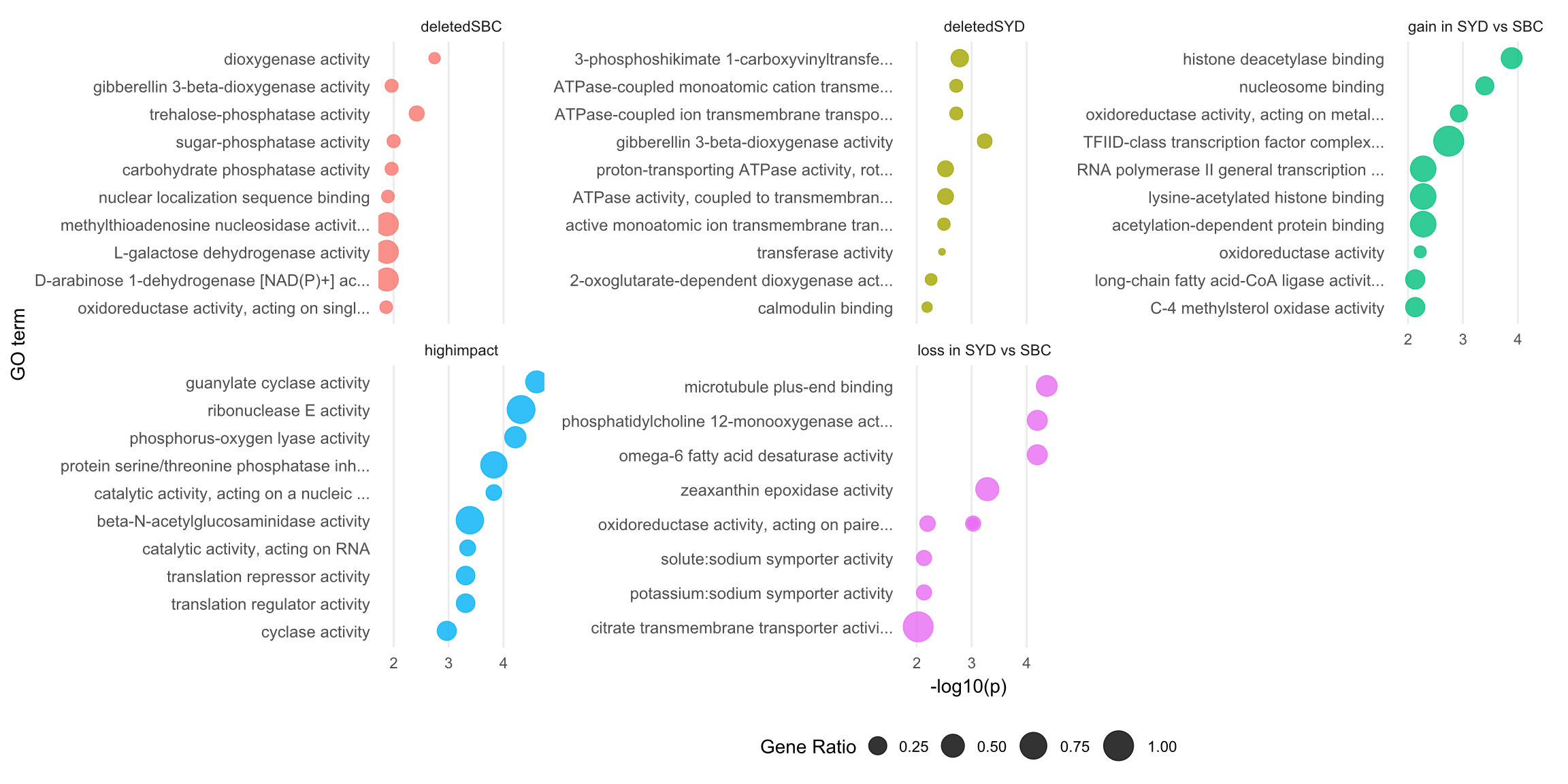

**Supplementary Figure 6:** GO terms enrichment in variable gene sets identified through comparison of diploid *Themeda triandra* accessions. Figures detail the significance of GO term enrichment in genes with CNV (gain or loss), putative deletions and genes with high impact mutations between a) PAN & SBC, b) PAN & SYD, c) SYD & SBC. The size of the bubble indicates what proportion of the entire set of genes annotated with that GO term are present in the CNV set. P-values are not adjusted for multiple testing.
